# Direct RNA Oxford Nanopore sequencing distinguishes between modifications in tRNA at the U_34_ position

**DOI:** 10.1101/2024.12.30.630739

**Authors:** W. Kusmirek, N. Strozynska, P. Martin-Arroyo Cerpa, A Dziergowska, G. Leszczynska, R. Nowak, M. Adamczyk

**Author notes:** first co-authors (W. Kusmirek, N. Strozynska).

## Abstract

The measurement of tRNA modifications with single transcript resolution has been feasible for only a few modifications due to the lack of available methods. This limitation does not allow to advance basic research studies on the dynamic nature of tRNA modification and its cellular function in time and space, neither to develop modern diagnostic tools for several already known tRNA-dependent human diseases. Nanopore is a well-established sequencing method that has proven to be efficient for the study of RNA. The analysis of tRNA modifications by Nanopore is still under development. We have investigated the efficacy of nanopore technology to discriminate between complex modifications of uridine 34 in tRNA, which affect the base-calling properties of neighbouring bases and are therefore difficult to accurately predict. We have developed new methods to chemically and enzymatically synthesise single modified tRNA molecules with modifications at the anticodon loop. Nanopore technology captures the features produced by uridine with and without a thiol group when present on synthetic tRNA molecules. Thus, Oxford Nanopore Technology (ONT) has great potential for developing strategies to accurately identify the modification status of the tRNA anticodon loop (ACL), which encompasses the most complex modifications on uridine-containing RNA motifs.

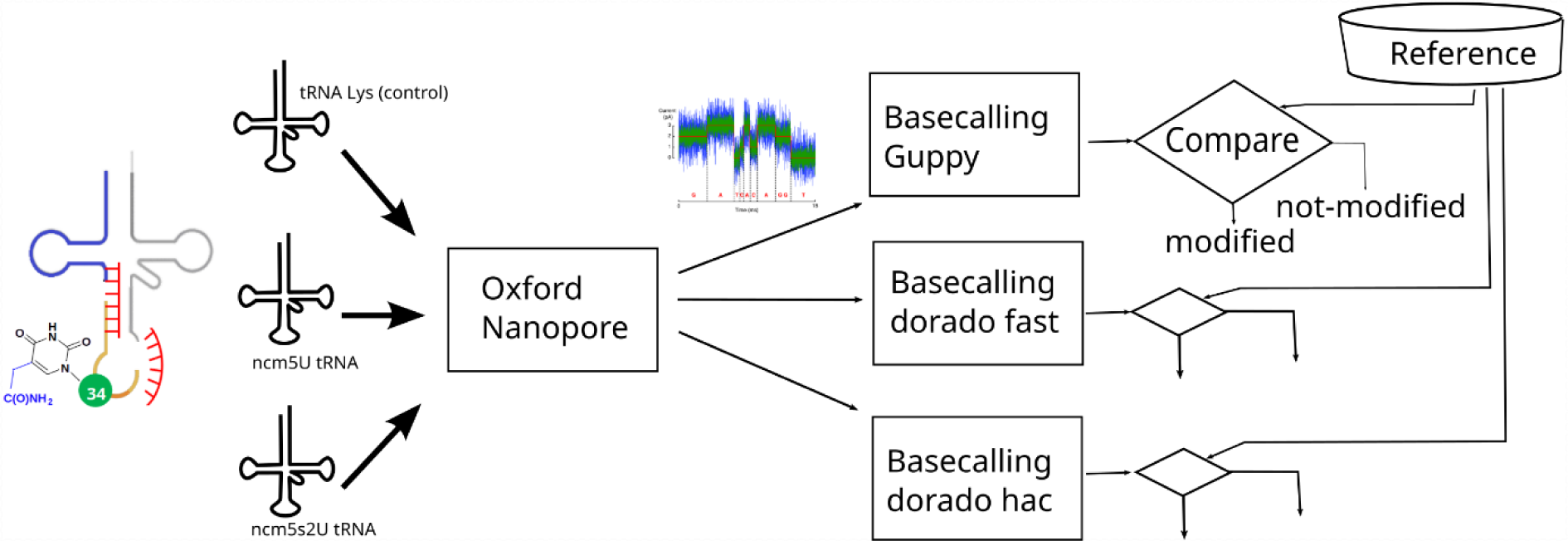

---

Transfer RNA (tRNA) is a molecule that is extensively modified post-transcriptionally by a variety of modifying enzymes, making this class of non-coding RNA the most diverse modification target among other types of RNA that perform different biological functions in cells. More than 100 modified nucleosides have been identified in tRNA species. Approximately 25 chemically distinct tRNA modifications can occur per tRNA molecule ^1^. In recent years, extensive research has uncovered associations between tRNA modification profiles and malignancy, primarily of neuronal systems, leading to neurological disorders. Not only the absence of tRNA modifications due to mutations in human modification genes or their downregulation is manifested in neurological and intellectual disability, but also in type 2 diabetes and tumorigenesis ^2–10^. There is a growing need for the development of modern, reliable and, above all, clinically accessible analytical methods for the detection of covalent tRNA modification disorders in the clinical setting.

tRNA modifications have various universal biological functions, facilitating correct tRNA folding and influencing translation capacity, efficiency and fidelity through the chemical modifications, interactions with tRNA molecules and specialised enzymes. There are currently over 70 enzymes that catalyse over 40 types of different tRNA chemical modifications. So far, tRNA modifications have been divided into two main functional groups according to their function, which correlates with the position of the nucleotide in the tRNA molecule ^11,12^. Modifications in the tRNA body from N1 to N30 and further from N40 to N76 influence tRNA stability and structure and play a role in tRNA interaction with ribosomes and synthetases. For example, pseudouridines add rigidity to the tRNA structure, dihydrouridines (D) facilitate tRNA elasticity, while N2, N2-dimethylguanosine (m^2,2^G) contributes to the two-dimensional structure of tRNA ^13,14^. Some of the modifications in this group are essential for establishing interactions of tRNA with modifying enzymes. The second group of modifications in the anticodon loop region includes positions from N31 to N39 in the anticodon arm. The ACL tRNA modifications play a role in modifying the process of mRNA decoding ^12^. The N34 and N37 modifications stabilise codon-anticodon interactions that have multiple A-U pairs and correctly discriminate between the wobble nucleotide (N34) and the third nucleotide of the codon (B3), whereas N37 enhances stacking interactions with the B1 base in the decoding centre of the ribosome A site ^15–17^. The universal modifications present in the three domains of life are those introduced by the Elongator Complex (ELP) and the Urm1 pathway on the uridine at position 34 of four cytosolic tRNAs; tRNA^Lys^_(UUU),_ tRNA^Gln^_(UUG),_ tRNA^Glu^_(UUC)_ and tRNA^Arg^_(UCU)._ ELP (Elp1, Elp2, Elp3 and Elp6) is responsible for the modification of uridine U_34_ to carboxymethyl uridine (xcm^5^U_34_) derived from acetyl-CoA, where x represents a methyl group (m) when esterified by Trm9 (human ALKBH8) and Trm112, or an amino group (n) ^18^. In addition, U_34_ can be further modified by thiolation, leading for example to the formation of 5-methoxycarbonylmethyl-2-thiouridine (mcm^5^S^2^U) by the Urm pathway activating thiol ^19^. In *S. cerevisiae* it has been shown that 5-carbamoylmethyluridine (ncm^5^U) and 5-carbamoylmethyl-2-thiouridine (ncm^5^S^2^U) are either the precursors of mcm^5^U (mcm^5^S^2^U) or the default modifications in the absence of the methyl modification ^20,21^. It has also been shown in yeast that the S^2^U_34_ modification is reduced in response to a shortage of sulphur amino acids, leading to reduced expression of the Uba4 protein (sulphur transferase) in the Urm pathway, suggesting that this modification is reversible under challenging nutritional conditions ^22,23^. Studies in haematopoietic stem cells (HSCs) suggest that mammalian cells may have a similar U_34_ modification-dependent sensing mechanism ^24^. The absence of U_34_ modifications (xcm^5^U or absence of the S^2^U moiety) is associated with multiple phenotypic effects such as temperature sensitivity, septate defects and proteotoxic stress, and can be rescued by overproduction of tRNA^Lys^_(UUU),_ tRNA^Gln^_(UUG)_ and tRNA^Glu^_(UUC)_ in model organisms ^21,25–27^. Mammalian phenotypes associated with impaired U_34_ modification have been mainly attributed to proteostasis, the stress-induced unfolded protein response (UPR) or impaired translation of specific proteins whose mRNA is enriched in the tRNA^Lys^_(UUU),_ tRNA^Gln^_(UUG)_ and tRNA^Glu^_(UUC)_ codons ^28,29^. In mammalian cells, the proteostasis imbalance was reversed by overexpression of unmodified tRNA, as in yeast cells ^30^.

As noted above, the lack of U_34_ modifications of many cytosolic tRNA targets and mitochondrial tRNA (mtRNA) leads to a confirmed variety of human neurodevelopmental disorders; familial dysautonomia, DEAM-PL syndrome, as well as cancer; melanoma, breast and colon cancer ^2,29,31,9,32,33^. In neuronal cell models and patients, tRNA U_34_ modifications also appear to be essential for maintaining brain function in the elderly ^30^. Studies in other cell types suggest that Elongator is required for tumour development or adaptation. On the contrary, the absence of the S^2^U_34_ moiety is associated with age-related clonal haematopoiesis, favouring the development of lymphoma, suggesting the multiple effects of U_34_ modifications on cellular homeostasis, which may be cell type dependent ^8^.

A detailed examination of the xcm^5^U and xcm^5^S^2^U modification status in cells can be performed quantitatively by liquid chromatography-tandem mass spectrometry (LC-MS/MS) at the nucleotide level. However, for the verification of "tRNA modopathies" in patients at single tRNA molecule resolution, sequencing-based methods seem to be more appropriate, offering simple solutions as a diagnostic tool independent of extensive laboratory infrastructure. The sequencing strategies for tRNA epitranscriptomics are under development. Currently, the direct RNA sequencing platform developed by Oxford Nanopore Technologies (ONT) allows the detection of a few modifications in native tRNA molecules ^34,35^. Computational algorithms allow the precise identification of several modifications including 6-methyladenosine (m^6^A), 7-methylguanosine (m^7^G), 5-methylcytidine (m^5^C), 5-hydroxymethylcytidine (hm^5^C), 5-formylcytidine (f^5^C), and pseudouridine (Ψ) in mRNA and rRNA using a nanopore direct RNA sequencing strategy and in vitro transcribed control synthetic RNA oligomers ^36,37^. EpiNano identifies m^6^A, although it cannot distinguish between different types of RNA modifications (e.g. m^1^A vs. m^6^A) ^38^. Algorithms for deciphering complex chemical modifications in tRNA are not yet established and offered in the ONT portfolio.

In this study, we aimed to map complex tRNA modifications such as ncm^5^U_34_ and ncm^5^S^2^U_34_ in tRNA^Lys^_(UUU)_ using Oxford Nanopore Sequencing (ONT) and to evaluate whether these types of RNA modifications can be associated with distinct signatures at the nanopore that could potentially be used to identify them. Using full length, synthetic and singly modified tRNA^Lys^, in this study we obtained ONT sequencing data for singly modified tRNA^Lys^_(UUU)_ placed at the wobble position U_34_ of the anticodon.

With this work, we contribute to the accurate identification of modifications at position U_34_ in the tRNA ACL. The current distribution curves observed for ncm^5^U_34_ and ncm^5^S^2^U_34_ indicate a measurable effect of the tRNA modifications on the nanopore signal patterns and distinguish the modifications. This promising result opens a new perspective for the detection of specific tRNA modifications at the U_34_ position in patients with neurological diseases, such as known dysautonomia and DREAM-PL syndrome, and metabolic disorders manifested by hypomodification of U_34_ in tRNA, by direct evaluation with advanced Oxford Nanopore Technologies.

## RESULTS AND DISCUSSION

### Design of the synthetic tRNA^Lys^_(UUU)_ and chemical synthesis of singly modified 13nt RNA oligomers

tRNA^Lys^ _(UUU)_ in *S. cerevisiae* has three modifications in the anticodon stem-loop at positions 34, 37 and 39, which are xcm^5^S^2^U_34_, mS^2^t^6^A_37_ and Ψ_39_. The uridine at position 34 is modified to either 5-carbonylmethyluridine (xcm^5^U_34_) or 5-carbonylmethyl-2-thiouridine (xcm^5^S^2^U_34_).

Studies by Durant et.al. 2005 ^39^ have shown that the modification of U_34_ has no effect on the structure of U_34_ or the ACL, while the S^2^U moiety promotes the stacking of U_34_ and U_35_ and increases the C3’-endo sugar conformation of U_34_. We wanted to verify whether the adopted conformation affects the sequence reading at the nanopore (ONT). Therefore, we constructed singly modified tRNAs using chemical and enzymatic approaches. To obtain the singly modified tRNAs specific for lysine, the enzymatic ligation of three chemically obtained RNA oligomers, including a 13 nt tRNA fragment containing one of four modified nucleosides at the first anticodon position. We selected two uridines and 2-thiouridines with methyl ester (abbreviated as mcm^5^U and mcm^5^S^2^U) or amide moiety (abbreviated as ncm^5^U and ncm^5^S^2^U) (Figure 1a). The synthetic work was planned in the following order: synthesis of the mcm^5^U and mcm^5^S^2^U building blocks of a 3’-*O*-phosphoramidite structure; synthesis of two ester-modified RNA oligomers by solid-phase phosphoramidite chemistry; deprotection of the oligomers in two ways, the first providing the residual ester and the second allowing efficient ester conversion to amide (Fig. 1c). Mcm5-uridine and 2-thiouridine nucleosides were synthesised in several steps by functionalising 5-bromouridine with an mcm5 substituent ^40^ or forming a *β*-*N*-glycosidic bond between silyl-activated mcm5-2-thiouracil and protected ribose ^41^. Both nucleosides were converted into the corresponding 3’-*O*-phosphoramidite building blocks (**1a, 1b**, Fig. 2) using standard protocols ^40,42^. Monomeric units **1a/1b** were used to synthesise two 13-nt RNA oligomers modified with ester containing mcm^5^U (**2a**) and mcm^5^S^2^U (**2b**). As the synthetic oligomers are fully protected and attached to the solid support (here abbreviated as CPG - controlled pore glass), the post-synthetic steps of oligomer deprotection were performed to release ‘free’ RNA. In addition, treatment of ester-containing oligomers with ammonia promoted ester ammonolysis, yielding amide-containing ncm^5^U and ncm^5^S^2^U-modified oligomers (**4a/4b**, Fig.2). To date, the conversion of mcm5 to ncm5 groups has only been reported for mcm^5^S^2^Uracyl and mcm^5^Uridine ^41^(41) and involved the treatment of ester-containing substrates with conc. ammonium hydroxide solution for 12 h at room temperature. In our protocol, the conc. ammonium hydroxide solution was replaced by a slightly heated mixture of conc. ammonia-ethanol in a ratio of 1:3. These conditions serve to efficiently remove exoamine masking groups and protect RNA from *tert*-butyldimethylsilyl (TBDMS) hydrolysis and subsequent RNA degradation ^43–45^.

**Figure 1:**
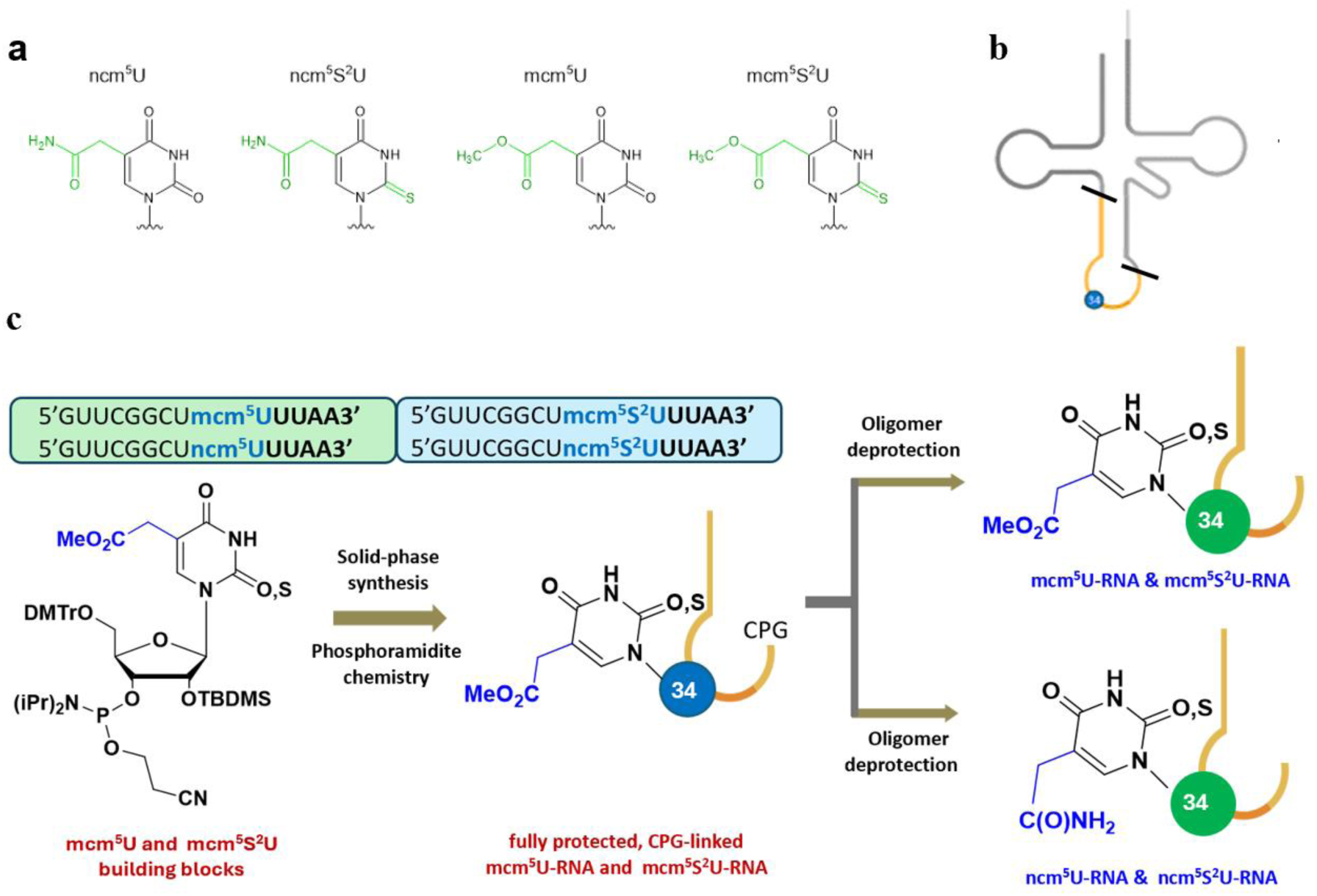
(a) Chemical structures of modified uridines (b) Schematic representation of singly modified tRNA at uridine 34 LEGEND: yellow loop indicates 13nt-modified RNA oligomer comprising ACL loop; grey loops indicate 25nt-RNA comprising D-loop and 38nt-RNA comprising T-loop (c) General strategy of 13nt-RNA synthesis. For a detailed description, see Materials and Methods.

**Figure 2.**
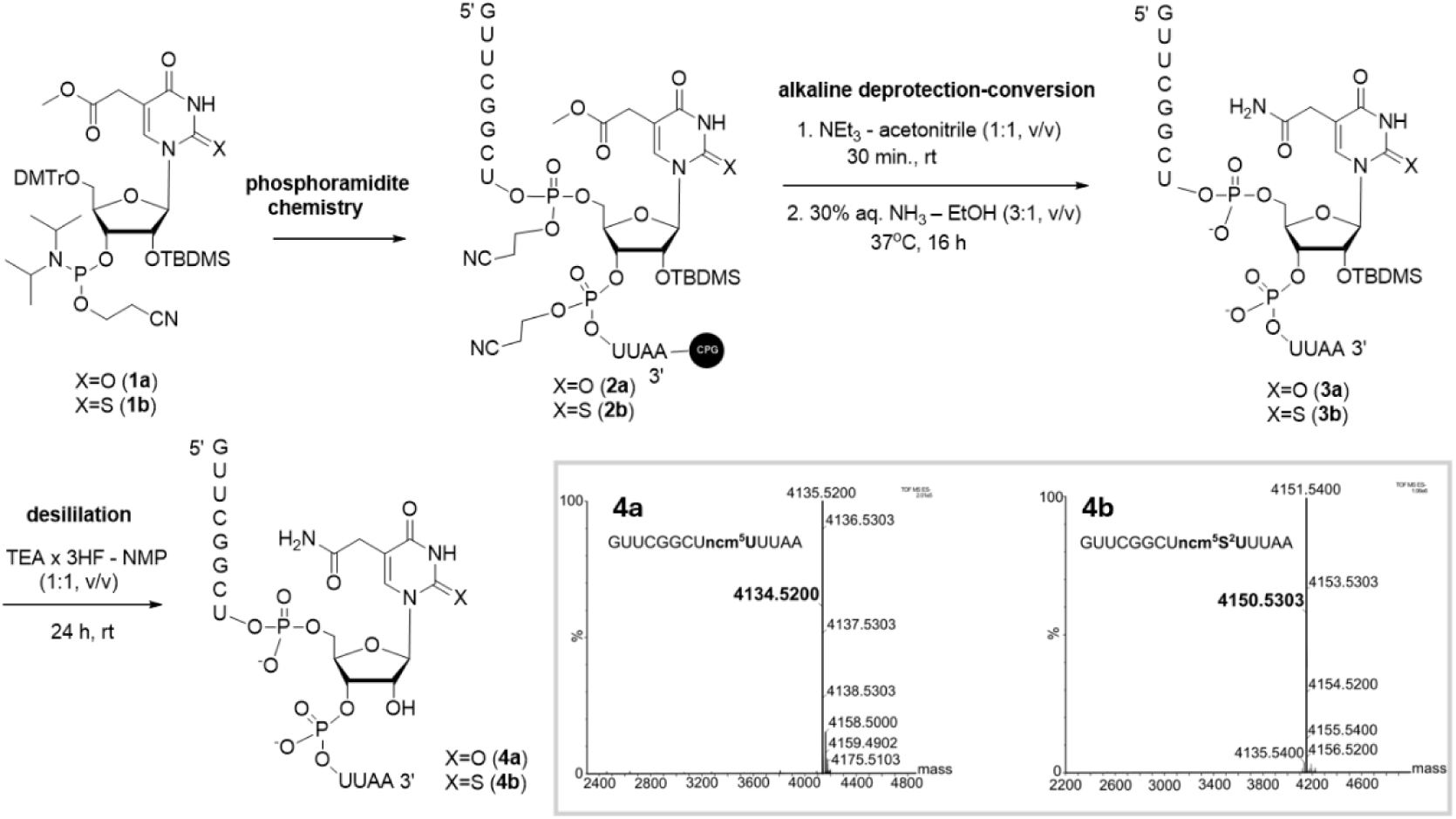
Scheme of preparation of mcm^5^U and mcm^5^S^2^U oligonucleotides **2a/2b** by phosphoramidite chemistry and further deprotection-conversion steps to give ncm^5^U and ncm^5^S^2^U RNA oligomers **4a/4b**. ESI MS spectrum of 5’GUUCGGCU**ncm^5^U**UUAA3’ (**4a**); calculated monoisotopic mass is 4134.52; measured m/z is 4134.51; ESI MS spectrum of 5’GUUCGGCU**ncm^5^S^2^U**UUAA3 oligonucleotide (**4b**); calculated monoisotopic mass is 4150.53; measured m/z is 4150.51. Electrospray mass spectrometry measurements were performed on a Synapt G2-Si mass spectrometer (Waters) equipped with a quadrupole time-of-flight mass analyser. The mass spectrometer was operated in negative ion detection mode. The results of the measurements were processed using the MassLynx 4.1 software (Waters) included with the instrument.

ESI MS spectrometry confirmed the correct structure of the modified oligomers and their purity (the calculated masses of the oligomers were identical to the measured masses (Figure 2).

Notably, the same ester-containing RNA oligomer **2a/2b** was effectively deprotected to yield free mcm^5^U RNA. To keep the ester moiety intact, a solution of DBU in anhydrous methanol was used in the alkaline RNA deprotection step to cleave the support and remove exoamine protecting groups ^15,46^.

### Optimisation of the split ligation reaction to obtain single modified full length tRNAs

Base calling errors and mismatch frequency can predict modifications carried by rRNA, tRNA and other non-coding RNAs in the direct RNA sequencing approach using Nanopore technology ^34,35,47^. Primary efforts have focused on accurately classifying nucleotides containing different modifications using synthetic RNA oligomers such as mRNA and rRNA to provide algorithms that accurately identify the modifications ^36–38^. Synthetic modified RNAs are typically obtained in IVT reactions. IVT-generated tRNA products occasionally suffer from limitations, such as the addition or omission of nucleotides at the 3’ end, and the reaction requires the use of a dsDNA template ^48^. Incorporation of a limited repertoire of commercially available modified nucleotides into IVT RNA may result in lower transcriptional efficiency compared to standard NTPs.

As an alternative, we designed universal 5’ and 3’ RNA oligos and ligated them by splint ligation methodology to 13nt RNA singly modified with uridine at position 34 of the ACL to assemble the full length tRNA^Lys^_(UUU)_ (Fig. 2a, Fig. S1).

We tested several different DNA splint oligos in the splint ligation reactions with the synthetic RNA oligomers to achieve the highest ligation efficiency. Only the best performing ones are reported in the supplementary material (Fig. S1, Table S1). The first ligation attempt was performed with an unmodified 13-nt RNA oligomer (at its 5’ end) and a 25-nt RNA fragment (Fig. S2a). As the UPLC MS chromatogram indicated incomplete ligation, the second experiment involved ligation between the 3’ ends of unmodified 13-nt and 38-mers.

A chromatogram of the oligomer mixture extracted after this ligation indicated complete ligation, yielding a correct 51-nt long product with a retention time of 8.36 minutes (Fig. 3b, Fig. S2a). The successful ligation of unmodified 13-nt and 38-nt fragments prompted us to use the developed conditions for the same type of ligation, but with modified ncm^5^U RNA. Again, complete ligation of the oligomers to the desired product was observed with a retention time of 6.52 and the correct mass (Fig. 3c). Initially, the ligation mixtures were resolved by UREA-PAGE, but the products did not migrate to the expected size (Fig. S2a, S2b). We successfully optimised the first and second ligation reactions by DNAse I treatment to remove DNA splint oligos, followed by gel extraction of the ligation products, which migrated as a single band. In all cases we confirmed the size of the 51 nt ligation product I and the 76 nt full length tRNA (ligation product II) by UREA-PAGE. The efficiency of ligation II reached 90% due to the prior 5’-end phosphorylation of the 25nt RNA oligomer. Following the optimised procedure, ncm^5^U_34_-tRNA^Lys^, ncm^5^S^2^U_34_-tRNA^Lys^, mcm^5^U_34_-tRNA^Lys^, mcm^5^S^2^U_34_-tRNA^Lys^ (Fig. 4b, 4d, 4f and 4h) and unmodified tRNA^Lys^ (Fig. S3) were generated and used for library construction and direct RNA sequencing. Our optimised protocols can be automated.

**Figure 3.**
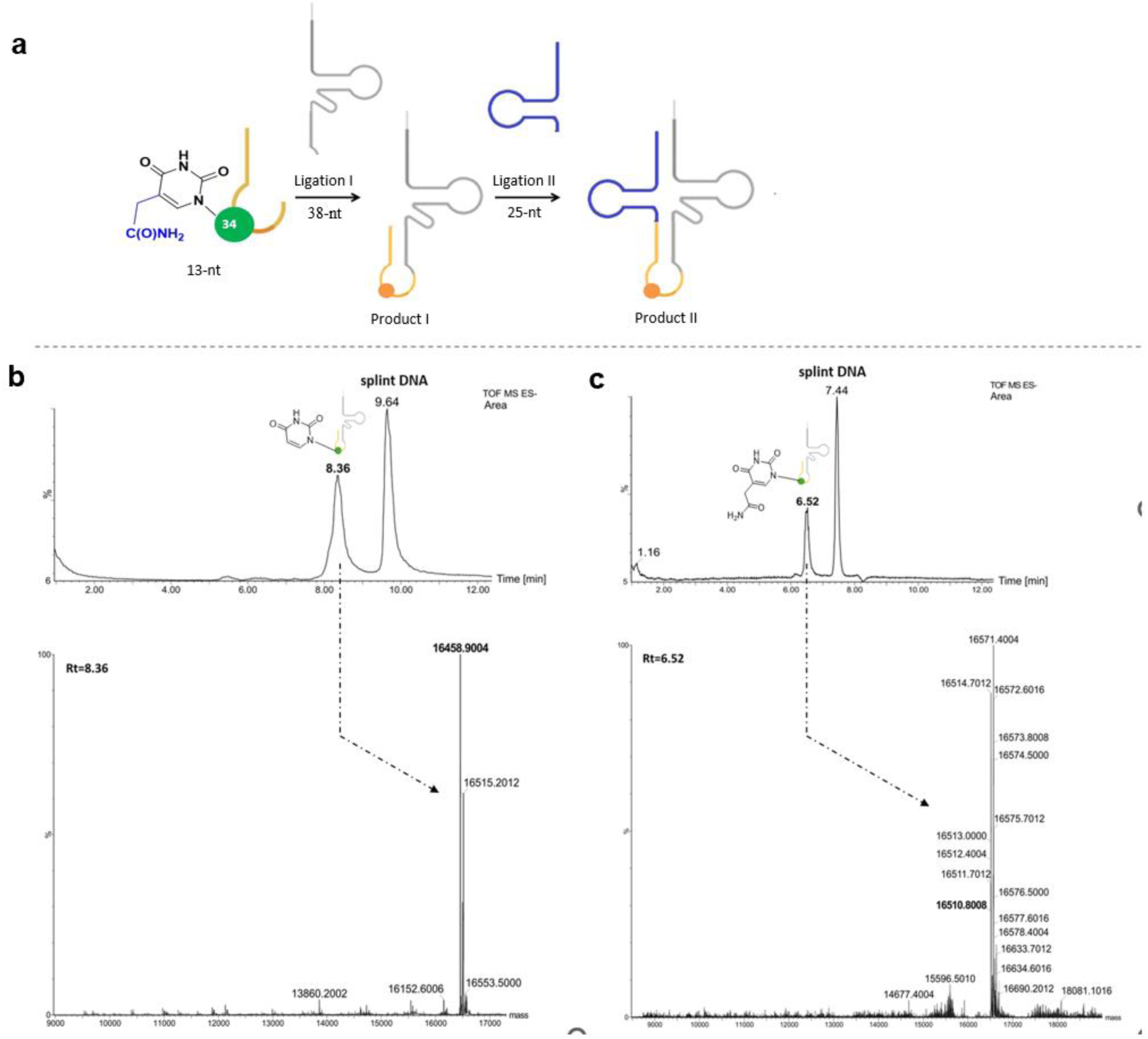
(a) Schematic of the general strategy for reconstitution of complete tRNAs by splint ligation using 13nt RNA, 38nt RNA (b-c) UPLC-PDA-ESI(-)-HRMS analysis of the 51nt product and deconvoluted mass spectra corresponding to (b) the unmodified 51nt ligation product with calculated monoisotopic mass 16458 (Fig. S2) or (c) the ncm^5^U-modified ligation product with calculated mass 16510 (Fig. S2).

**Figure 4.**
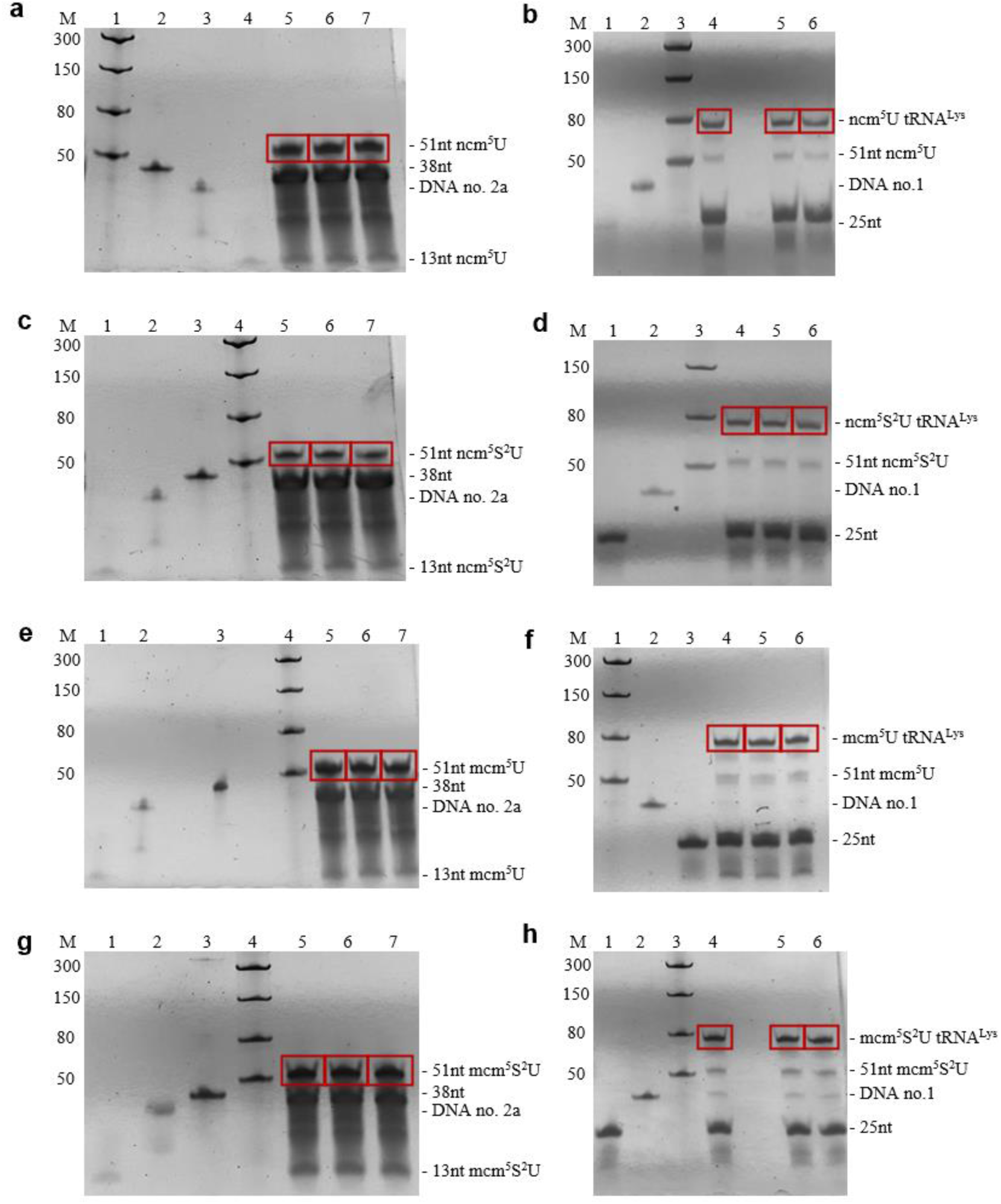
UREA-PAGE of the ligation reaction mixtures after DNase I treatment. (a) Ligation I: Line 1: Low Range ssRNA ladder; Line 2: 38nt RNA; Line 3: DNA No.2a; Line 4: 13nt ncm^5^U RNA; Lines 5-7: 51nt ncm^5^U-RNA (b) Ligation II: Line 1: 25nt RNA; Line 2: DNA No.1; Line 3: Low Range ssRNA Ladder; Lines 4-6 ncm^5^U_34_-tRNA^Lys^. (c) Ligation I: Line 1: 13nt ncm^5^S^2^U-RNA; Line 2: DNA no. 2a; line 3: 38nt; Line 4: Low Range ssRNA Ladder; Lines 5-7: 51nt ncm^5^S^2^U-RNA; (d) Ligation II: Line 1: 25nt RNA; Line 2: DNA No.1; Line 3: Low Range ssRNA ladder; lines 4-6: ncm^5^S^2^U_34_-tRNA (e) Ligation I: line 1: 13nt mcm^5^U-RNA, line 2: DNA no. 2a; line 3: 38nt RNA; Line 4: Low Range ssRNA ladder; Lines 5-7: 51nt mcm^5^U-RNA (f) Ligation II: Line 1: Low Range ssRNA ladder; Line 2: DNA No.1; Line 3: 25nt RNA; Lines 4-6: mcm^5^U_34_-RNA^Lys^ (g) Ligation I: 13nt mcm^5^S^2^U-RNA; Line 2: DNA no. 2a; Line 3: 38nt RNA; Line 4: Low Range ssRNA Ladder; Lines 5-7: 51nt mcm^5^S^2^U-RNA (h) Ligation II: Line 1: 25nt RNA; Line 2: DNA No.1; line 3: Low Range ssRNA ladder; lines 4-6: mcm^5^S^2^U_34_-tRNA^Lys^. The unmodified and modified 51nt RNA and full tRNA^Lys^ ligation product bands are highlighted in red.

### ONT library preparation using tRNA does not require a reverse transcription step

The direct RNA sequencing method developed by Nanopore MinION is particularly useful for sequencing mature tRNA strands. Early efforts to sequence tRNA molecules have demonstrated the reliability of the sequencing strategy, which does not include the reverse transcription (RT) step in the library construction procedure typically performed and recommended for mRNA sequencing by ONT ^34^. We adopted the approach of Thomas et al. 2021 with slight modifications to allow detection of full-length tRNA^Lys^ through 5’ and 3’ extensions using custom splint adapters and omitting the RT step ^34^. Hybrid RNA/DNA double-stranded adapters were ligated to the NCCA 3’ tRNA end and 5’ termini using T4 RNA Ligase 2 and run on 8% denaturing PAGE gels (Fig. 5a, 5b, Fig. S4). We used a 20-fold higher concentration of the splint double-strand adapter than tRNA. The improvement of the efficiency of tRNA ligation with adapters to 60-70% was dependent on the annealing of the splint adapters and on the prior 5’ phosphorylation of the tRNAs (Materials and Methods). Resulting ligation products in the range of 100-140 nt were extracted from gels and purified. In all ligation mixtures visualised by UREA-PAGE, we obtained two ligation products (indicated by red boxes). The purified, modified and unmodified tRNAs were ligated to the ONT adaptor provided in the Direct RNA Sequencing Kit (SQK-RNA002) using T4 DNA ligase. The presence of modifications on synthetic RNA reduces read efficiency.

**Figure 5.**
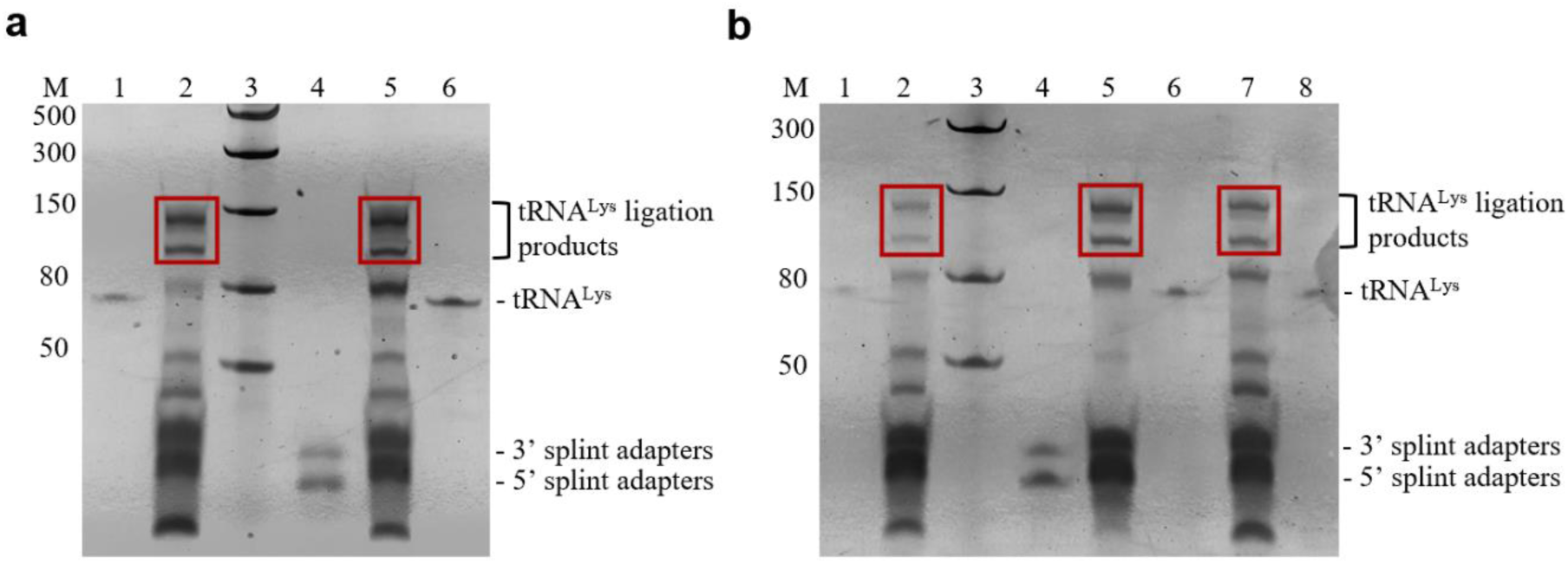
UREA-PAGE of ligation reaction mixtures of tRNA^Lys^ with adapters. (a) Line 1: ncm^5^S^2^U_34_-tRNA^Lys^; Line 2: ligation products of ncm^5^S^2^U_34_-tRNA^Lys^ with adapters; Line 3: Low Range ssRNA ladder; Line 4: 5’ splint and 3’ splint adaptors; line 5: ligation products of unmodified tRNA^Lys^ with adaptors; line 6: unmodified tRNA^Lys^ (b) Line 1: ncm^5^U_34_-tRNA^Lys^; line 2: ligation products of ncm^5^U_34_-tRNA^Lys^ with adaptors; line 3: Low Range ssRNA ladder; Line 4: 5’ splint and 3’ splint adapters; Line 5: ligation products of mcm^5^S^2^U_34_-tRNA^Lys^; Line 6: mcm^5^S^2^U_34_-tRNA^Lys^; Line 7: ligation products of mcm^5^U_34_-tRNA^Lys^; Line 8: mcm^5^U_34_-tRNA^Lys^. The ligation products are shown in red.

### A basecaller significantly affects the ability to detect tRNA modifications

Numerous artificial intelligence tools and models have been developed specifically for the basecalling process. Previous work has shown that basecalling errors or mismatches to the reference can be used to detect RNA modifications ^35,37,49–53^. In agreement with a report by White et al. ^54^, complex modifications of uridines were observed as base calling errors in the sequenced tRNA^Lys^_(UUU)_ in this study. Our results highlight the significant impact of the selected basecalling tool and model on the detection of tRNA modifications, as data from different tools and models resulted in different read alignments to the reference tRNA sequence. To identify the optimal basecaller for the detection of tRNA modifications at U_34_, we used the Guppy and Dorado basecallers using both the fast and hac models (Table S2 and Table S3). Visualisation of the reads using the IGV tool showed that for all three sets of reads (from the Dorado basecaller with fast and hac models and from the Guppy basecaller) there are visible mismatch errors in the ncm^5^U_34_ and ncm^5^S^2^U_34_-tRNA^Lys^ modification region (Figure 6a). However, it was found that different basecallers report different levels of base modifications for uniformly modified synthetic tRNA (Figure 6b). This is particularly evident in the ratio of uracil to cytosine levels for the ncm^5^U_34_ and ncm^5^S^2^U_34_ modification error signatures. Consistent with previous observations on biological tRNA, the modifications present at position U_34_ of tRNA^Lys^ caused changes in the level of U-to-C mismatches, which clustered in the vicinity of U_34_ within 1-2 nucleotides of the modified base ^54^. This is related to the fact that the Oxford Nanopore sequencer analyses the current value for a window of a few bases, but rarely for complex modifications at single base resolution. However, using Dorado hac and fast, we observed the highest levels of U-to-C mismatches at the U_34_ and U_35_ positions. Despite the general differences in ribonucleotide A, C, G and U levels reported by the software, we observed that there is a higher level of U-to-A mismatch detection in base called features for the ncm^5^S^2^U_34_ modification compared to the ncm^5^U_34_ modification without thiolation (Fig. 6b).

**Figure. 6.**
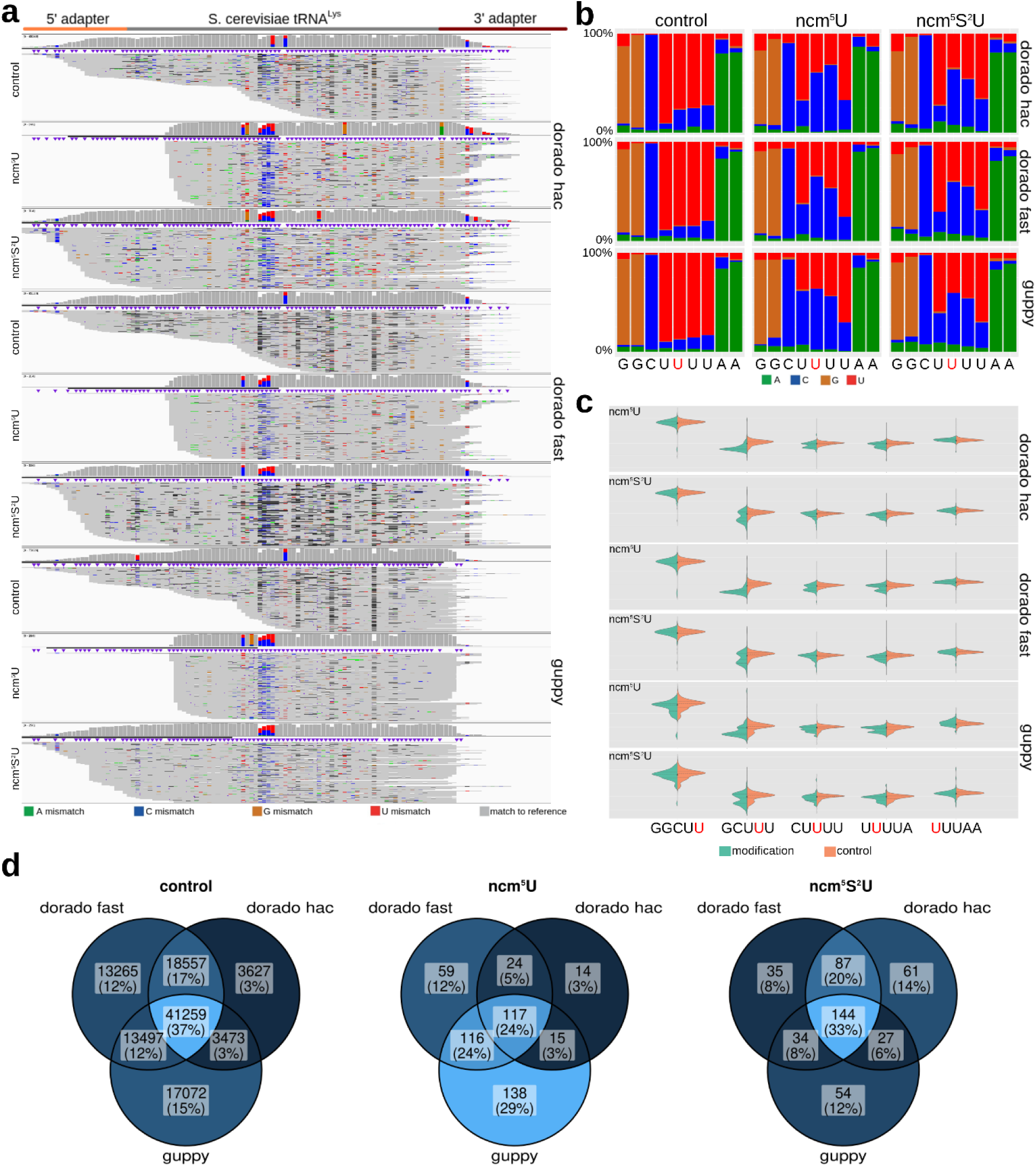
The choice of basecaller significantly affects the ability to detect tRNA modifications. (a) IGV application snapshot of reads generated by the Dorado basecaller (fast and hac models) and by the Guppy application mapped to tRNA^Lys^. The tRNA^Lys^ reference sequence prior to mapping was extended with adapter sequences marked in orange (5’ splint adapter) and red (3’ splint adapter). (b) Distribution of RNA bases in reads generated by different basecallers in the region of the studied modification. (c) Effect of different basecallers on the current values in the tRNA modification region. The signal is generated for each k-mer and represents a nucleotide at position 1 in k-mer (d) Venn diagram showing the number of reads mapped to a reference.

Since different basecallers generate different sets of tRNA reads, once the reads are mapped to the reference sequence, a different subset of reads (and their associated currents) defines the distribution of currents in the modification region (Fig. 6c). Notably, the Dorado basecaller shows the highest current signal bias with a shift of −3 nt from the modified U in both tRNA^Lys^ (Fig. 6c see GCUUU motif). Instead, using the Guppy tool results in an additional signal anomaly for ncm^5^U in the GGCUU k-mer where the modified uridine is placed at the 3’ end (Fig. 6c). These observations are crucial for downstream applications and for building AI models to detect tRNA U_34_ modifications, as a model trained on data from a base caller should be used and tested with the same base caller. Its effectiveness will be significantly reduced with a different basecaller.

We investigated how many reads from different basecalling methods and models lead to the recovery of tRNA that maps to the reference sequence (Fig. 6d). We found that, across the three datasets, only a small subset of reads overlaps between the different basecalling methods. Our results emphasize that using different basecallers can sometimes produce different sequences. This means that it is worth analysing the same raw data sets using different basecallers to find uncertain elements.

### Current methodologies demonstrate efficacy in identifying ncm^5^U_34_ and ncm^5^S^2^U_34_ tRNA^Lys^ modifications on synthetic data

Several tools and methodologies have been developed to detect tRNA modifications. We investigated how existing tools would perform on data generated by ncm^5^U_34_-tRNA^Lys^ and ncm^5^S^2^U_34_-tRNA^Lys^ sequencing. We used the Nano-tRANseq ^35^, nanoCEM ^55^ and Nanocompore ^50^ tools. The Nano-tRNAseq tool correctly detects the modifications present on tRNA^Lys^ with minor differences (Fig. 7a, Fig. S5a, Fig. S6a). This tool does not use current values to detect modifications, but instead calculates the probability of a modification occurring based on the number of errors mapping reads compared to the reference sequence. Next, we tested two applications that use RNA reads and raw current values as an average to detect and determine the location of modifications in tRNA. NanoCEM consistently failed to correctly detect the ncm^5^U_34_ modification in all data tested. On the other hand, this tool detects the ncm^5^S^2^U_34_ modification at U_35_ with a shift of +1nt, as shown in the MANOVA distribution plot (Fig. 7b). However, the highest value is read for position G30 in k-mer GGCUU, which represents a −4 nt shift from modified U_34_. The best result was obtained using the Nanocompore tool, which illustrates the different effects of nucleotide modification for the two variants shown in Fig. 7, panel c. In the case of both modifications, the GMM and KS_intensity −log(p-values) reaches the maximum value at U_33_, representing k-mer GCUUU. Notably, the GMM and KS_intensity log(p-values) at position U_36_ shows a measurable effect of the ncm^5^U_34_ vs. ncm^5^S^2^U_34_ modification in k-mer UUUUUA.

**Figure 7.**
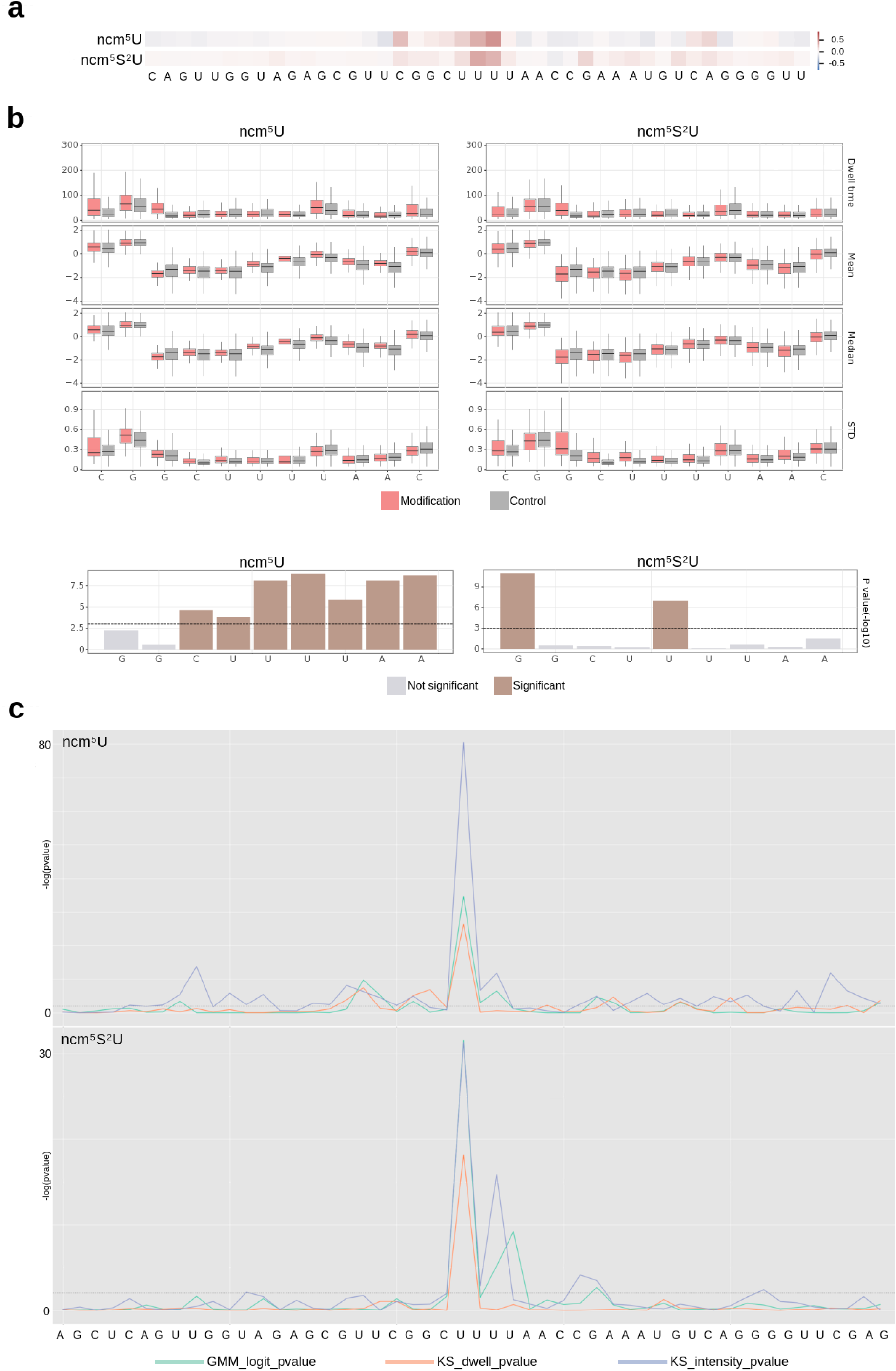
Detection of ncm^5^U_34_-tRNA^Lys^ and ncm^5^S^2^U_34_-tRNA^Lys^ using nano-tRNAseq, nanoCEM and Nanocompore tools. Detection of modifications of ncm^5^U_34_ and ncm^5^S^2^U_34_-tRNA^Lys^ using Nano-tRNAseq, nanoCEM and Nanocompore tools. (a) Heatmap of summed base calling error frequency by Nano-tRNAseq. (b) The top panel shows box plots describing the distribution of dwell time, mean, median and standard deviation of current for all RNA reads. The bottom panel shows the position where there is the greatest difference for the modified sample compared to the control, according to the nanoCEM application. (c) P-values for three algorithms comparing the modified sample with the control sample provided by Nanocompore applications: bivariate classification method based on 2-component Gaussian mixture model clustering followed by logistic regression test (GMM_logit) and robust univariate pairwise tests (Kolmogorov-Smirnov test) on dwell time (KS_dwell) or current intensity (KS_intensity).

### The mapping algorithm and its parameters significantly affect the ability to detect tRNA modifications

We observed that the choice of algorithm for mapping reads to the reference sequence, as well as the associated parameter settings, significantly influenced the ability to detect tRNA modifications. First, we examined how different algorithms and parameter configurations affected the number of correctly mapped reads (Tables S5-S7). The results showed that for all three basecallers, the minimap2 algorithm mapped a significantly lower number of reads compared to the other methods. Specifically, the dorado basecaller using the rna002_70bps_hac@v3 model for the control sample produced the highest number of tRNA reads (10.01% of the total mapped tRNA reads) when mapped with minimap2. In contrast, minimap2 consistently mapped less than 10% of the tRNA reads for the remaining samples and basecallers.

Second, we observed that different mapping algorithms and their parameter sets affected the mapping quality, as indicated by the bar graphs showing the alignment identity for different conditions in Figs. S7, S8 and S9 in panels E. These graphs show that minimap2 achieved the highest alignment identity; however, as shown in panels D, this was at the cost of a very low number of mapped reads. Additionally, when comparing the results of the bwa mem algorithm with different -T parameter values (bwa mem -W13 -k6 -xont2d -T30, -T20 and -T10), we found that as the value of T decreased, the number of mapped reads increased but the alignment identity decreased. This trend was consistent across all samples and base callers.

Finally, we evaluated the number of reads that were mapped to the reverse (antisense) strand. The data obtained show that the largest number of incorrect reverse strand mappings occurred with the bwa sw -z10 -a2 -b1 -q2 -r1 parameter set (Fig. S7f, Fig. S8f, and Fig. S9f).By comparing the number of uniquely aligned reads and the incorrect alignments in panels D, E, and F in Fig. S7, Fig. S8 and Fig. S9, we concluded that the bwa mem algorithm with the parameter set -W13 -k6 -xont2d -T30 is the most effective for mapping tRNA reads to the reference sequence.

## CONCLUSIONS

In this study, we have developed an efficient semi enzymatic method for the rapid production of modified tRNAs. The method can be used to assemble a variety of tRNAs by using custom oligomers and, *ad libitum*, modified oligomers containing site-specific incorporations of modified nucleosides. In addition, a facile and efficient protocol for post-synthetic ammonolysis of chemically obtained mcm^5^(S^2^)U-modified 13nt oligomers was elaborated yielding ncm^5^(S^2^)U-RNA counterparts.

We used the full length singly modified tRNAs to collect sequencing data. The sequencing data obtained in this study confirm that the ONT pore with two reading heads (R10.4 Chemistry) provides the accuracy and resolution of homopolymeric regions to capture the structural difference in linearised, single strand tRNA modified by ncm^5^U_34_ and ncm^5^S^2^U_34_.

Moreover, the choice of the basecaller (Dorado Hac, Dorado Fast, or Guppy) significantly affects the ability to detect tRNA modifications.

The aberrant specific modification of uridine 34 in tRNA has been widely recognised in human disease. The results of this study offer a starting point for the future development of a diagnostic tool to assess U_34_ modifications in biological tRNAs, which will be fully based on Oxford Nanopore technology.

## MATERIALS AND METHODS

### 13-nt RNA oligomers modified with 5-carbamoylmethyluridine (ncm^5^U) and 5-carbamoylmethyl-2-thiouridine (ncm^5^s^2^U)

Oligonucleotides with the sequences 5’GUUCGGCU**ncm^5^U**UUAA3’ and 5’GUUCGGCU**ncm^5^S^2^U**UUAA3’ (**4a/4b**, Figure 2) were prepared by post-synthetic conversion of the precursor 5’GUUCGGCU**mcm^5^U**UUAA3’ and 5’GUUCGGCU**mcm^5^S^2^U**UUAA3’ oligomers (**2a/2b**), respectively. The synthesis of the **2a/2b** precursors was performed by solid phase *β*-cyanoethyl phosphoramidite chemistry on a CPG support according to protocols described in the literature ^15,40,46^, using 5’-*O*-dimethoxytrityl-2’-*O*-*tert*-butyldimethylsilyl protected mcm^5^U and mcm^5^S^2^U phosphoramidites **1a/1b** as monomeric units ^40–42^.

After assembly, the support-linked DMTr-off oligomer **2a/2b** containing mcm^5^U or mcm^5^S^2^U was subjected to the deprotection-conversion steps on a 0.2 μmol scale. The *β*-cyanoethyl groups were selectively removed from the phosphate residues using Et_3_N-acetonitrile (260 μL, 1/1, v/v, 30 min, rt.). The supernatant was removed, and the CPG was washed twice with dry acetonitrile. The resin free of *β*-acrylonitrile was dried and treated with a 3:1 mixture of 30% NH_4_OH and ethanol (300 μL) for 16 h at 37 °C to cleave the nucleobase protecting groups and the CPG solid support. Simultaneously with these reactions, ammonolysis of the ester residues in mcm^5^U/mcm^5^S^2^U RNAs was performed, yielding ncm^5^U/ncm^5^S^2^U amide counterparts **3a/3b**.

After the deprotection-conversion step, the supernatant containing partially deprotected ncm^5^U/ncm^5^S^2^U oligomer **3a/3b** was transferred to an Eppendorf tube. The solid support was washed twice with a mixture of EtOH-H_2_O (1:1, v/v). The combined washes were dried on a Speed-Vac. To remove the TBDMS groups, each oligomer **3a/3b** was dissolved in 1-methyl-2-pyrrolidinone (NMP, 60 μL) and incubated with triethylamine trihydrofluoride (Et_3_N-3HF, 60 μL) for 24 h at room temperature. Fully deprotected RNA **4a/4b** was precipitated with ethoxytrimethylsilane, dried under vacuum and purified by anion-exchange HPLC on a Source 15Q 4.6/100PE column using a gradient of sodium bromide in a phosphate buffer containing 50 µM ethylenediaminetetraacetic acid. The oligomer-containing fraction was desalted on a C-18 cartridge (Sep-Pak^®^, Waters). After evaporation, ncm^5^U and ncm^5^S^2^U oligomers **4a/4b** were characterised by ESI MS (Fig. 2).

### Ligation of RNA oligomers by splint ligation

Synthetic RNA oligos were provided by Sigma Aldrich UK (Fig. S1). The singly modified 13nt-RNA was 5’ phosphorylated using T4 PNK (10U/µl New England Biolabs) according to the protocol provided by the manufacturer. Singly modified 13nt-RNA (1nmol) and unmodified 13nt-RNA (1nmol) were first ligated with 38nt-RNA (1nmol) in the presence of splint oligo DNA no. 2a (0.5nmol) (Table S1), PEG4000 (Thermo Scientific), ATP (Thermo Scientific) and T4 DNA ligase (5U/µl Thermo Scientific) in a total reaction volume of 500µl ^56^. The reaction mixtures were incubated at 37°C for 2 h. DNase treatment of the final ligation reaction mixtures with DNase I RNase-free enzyme (1 U/µl Thermo Scientific) was performed prior to UREA-PAGE. Samples were purified using the Monarch RNA Cleanup Kit (New England BioLabs). The substrates and products (51-nt RNA oligomers) of the reactions were separated on UREA-PAGE (acrylamide/bisacrylamide (29:1) (Sigma-Aldrich). The efficiency of the ligation reaction was 90%. After purification of the bands representing 51nt-RNA from the polyacrylamide gel, their MW was verified by LC-MS. 51nt RNA oligomers were further used in ligation reactions with 25nt RNA (0.1nmol), splint oligo DNA No.1 (0.05nmol) in a total reaction volume of 50µl (in the presence of PEG4000, ATP and T4 DNA Ligase Thermo Scientific). The 25nt RNA was additionally 5’-phosphorylated (T4 PNK 10U/µl NEB) in a total reaction volume of 50µl. Samples were incubated at 37°C for 2 h. The efficiency of the reaction, which yielded 76nt RNA, was lower and estimated to be 70%. The efficiency of the reaction was improved to 90%. The final products of the splint ligations were visualised on 8% UREA-PAGE using SYBR Gold nucleic acid gel stain (Invitrogen). For prolonged storage, individually modified and unmodified tRNA^Lys^ were precipitated with propan-2-ol in the presence of glycogen (20 mg/ml Thermo Scientific) and stored at −80°C.

### Ultra-performance liquid chromatography coupled with mass spectrometry and photodiode array detection (UPLC-PDA-ESI-MS)

The efficient splinted ligation of the 13-nt oligomer (unmodified or modified with ncm^5^U) with the 38-nt tRNA fragment (Fig. S2a and S2b) was confirmed by the UPLC-PDA-ESI(-)-MS method using an ACQUITY UPLC I class chromatography system equipped with a photodiode array detector with a binary solvent manager (Waters Corp, Milford, MA, USA) coupled to a SYNAPT G2-Si mass spectrometer equipped with an electrospray source and a quadrupole time-of-flight mass analyser (Waters Corp., Milford, MA, USA). An ACQUITY UPLC® Oligonucleotides BEH C18 column (50 × 2.1 mm, 1.7 µm, 60^◦^C) was used for the chromatographic separation of oligonucleotides. A gradient programme was used for the mobile phase, combining solvent A (15 mM triethylamine and 400 mM 1,1,1,3,3,3-hexafluoropropan-2-ol in water) and solvent B (50% methanol and 50% solvent A, v/v) as follows 33-37% B (0-1.0 min), 37-47% B (1.0-10.0 min), 47% B (10.0-10.5 min), 47-33% B (10.5-10.6 min) and 33% B (10.6-12 min). The flow rate was 0.2 mL/min (Fig. 2A) or 0.3 mL/min (Fig. 2b) and the injection volume was 10 µL. For mass spectrometric detection, the electrospray source was operated in negative resolution mode. The optimised source parameters were: capillary voltage, 2.7 kV; cone voltage, 40 V; desolvation gas flow rate, 600 L/h; temperature, 400^◦^C; nebuliser gas pressure, 6.5 bar; source temperature, 120 ^◦^C. Mass spectra were recorded over the range m/z 400 to 2000. The system was controlled by MassLynx v 4.1 software. The raw ESI mass spectra were deconvoluted to a zero charge state mass using the MaxEnt1 algorithm.

### Direct preparation of tRNA^Lys^ libraries

Full-length tRNAs (76nt-RNAs), unmodified and singly modified, were ligated with universal adapters and adapter oligo GCCA Lys (Table S1) according to Thomas et al 2021 ^34^ with some minor modifications. 5µM stocks of double-stranded adapters were prepared in MiliQ water by adding 100 pmol of each strand in a total volume of 20µl. The oligos were hybridised at 75°C for 15s and slowly cooled to 37°C. The reaction mixtures contained adapters, ATP (Thermo Scientific), PEG 4000 (Thermo Scientific), T4 RNA Ligase 2 (10U/µl New England Biolabs) and were incubated at 37°C for 2 h. Samples were purified using the Monarch RNA Cleanup Kit (New England Biolabs) and transferred to fresh, low-bind Eppendorf tubes. Library preparation was performed according to the Direct RNA Sequencing protocol (SQK-RNA002). The ligation reaction mixtures contained tRNA^Lys^ extended with double-stranded adapter oligo, NEBNext Quick ligation reaction buffer, RNA adapter (RMX), T4 DNA Ligase (2000 U/µl New England Biolabs) in a total volume of 25µl. The reactions were treated with RNA Clean XP beads (1.8x) at RT for 15 min. The RNA bound to the magnetic beads was then washed with Wash Buffer (WSB) on a magnetic rack (Promega). RNA was eluted with Elution Buffer (EB) after 20 min incubation at RT and prepared according to the original Direct RNA Sequencing protocol for loading onto the primed MinION flow cell.

### Sequencing conditions using the MinION flow cell

Sequencing was performed on a laptop running Windows 10, MinKNOW software ver. 23.07.15, MinKNOW Core ver. 5.7.5. The kit SQK-RNA002 was used, and the run was limited to 72 hours with active channel selection, pore reservation and pore scan frequency of 1.5 hours. Base calling was disabled. Data output was set to FAST5 and Bulk file; FASTQ and BAM output were disabled.

### Bioinformatic analysis

Basecalling was performed on the fast5 raw reads using the Guppy (ver. 5.0.7) and Dorado (ver. 0.4.1) applications; https://github.com/nanoporetech/dorado; 2024. The Guppy basecaller was started in high accuracy mode using the guppy_rna_r9.4.1_70bps_hac model, while the Dorado basecaller was started in two modes: fast and high accuracy using the rna002_70bps_fast@v3 and rna002_70bps_hac@v3 models, respectively. The statistics of the reads obtained are shown in Supplementary Table S3. Base-called reads were filtered for length and quality using the tools BBMap ^57^ (ver. 38.73) and fastq-filter https://github.com/LUMC/fastq-filter (ver. 0.3.0). We removed reads longer than 200 bp and with a mean base call accuracy of less than 80% (fastq-filter -q 7 -L 200) from further analysis (Table S4). We then generated reference sequences - we downloaded a set of unmodified S. cerevisiae RNAs from MODOMICS ^1,58^ and extended each RNA by adding the sequence of 5’ and 3’ splint adapters. We then mapped the resulting sets of base-called reads to the prepared reference sequence using the minimap2 ^59^ (ver. 2.17-r941) and BWA ^60^ (ver. 0.7.17-r1188) applications. The following parameter configurations were tested: (i) minimap2 -ax map-ont -k15; (ii) bwa mem -W13 -k6 -xont2d - T30; (iii) bwa mem -W13 -k6 -xont2d -T20; (iv) bwa mem -W13 -k6 -xont2d -T10; (v) bwa mem -W9 -k5 -xont2d -T10; (vi) bwa bwasw -z10 -a2 -b1 -q2 -r1. The best parameter set (bwa mem -W13 -k6 -xont2d -T30) was selected based on the number of uniquely mapped reads and reads mapping to antisense strands (mismapping) (Table S5, S6 and S7). Alignment identity was then quantified using SAMtools stats ^60^ (ver. 1.10), and the resulting bam files were visualised using the IGV software tool ^61–64^. After visualisation of the bam files, we start the previous applications for detection of tRNA modifications: Nano-tRNAseq ^35^, nanoCEM ^55^ and Nanocompore ^50^.

The scripts used for basecalling, quality control, mapping and subsequent bioinformatic analyses are available online at: https://github.com/wkusmirek/nanopore-tRNA-analysis.

## Acknowledgements

We would like to thank Katarzyna Pietka for her excellent technical support. This project was funded by the Excellence Initiative: Research University (IDUB) programme. Project BIOTECHMED-3 Advanced under grant agreement number 1820/334/Z01/POB4/2021 to M.A.

## Authors contributions

Conceptualization, M.A.; methodology, M.A., G.L. and R.N.; formal analysis, M.A., N.S., D.A., M-A.C.P and W.K.; investigation, M.A. G.L. W.K. and R.N.; resources, M.A.; data curation, M.A., G.L., W.K.; writing—original draft preparation, M.A.; writing—review and editing, M.A., G.L, W.K.; visualization, M.A., N.S., G.L., W.K and R.N.; supervision, M.A., project administration, M.A.; funding acquisition, M.A.. All authors have read and agreed to the published version of the manuscript.

## REFERENCES

(1) Boccaletto, P.; Stefaniak, F.; Ray, A.; Cappannini, A.; Mukherjee, S.; Purta, E.; Kurkowska, M.; Shirvanizadeh, N.; Destefanis, E.; Groza, P.; Avşar, G.; Romitelli, A.; Pir, P.; Dassi, E.; Conticello, S. G.; Aguilo, F.; Bujnicki, J. M. MODOMICS: A Database of RNA Modification Pathways. 2021 Update. Nucleic Acids Res 2022, 50 (D1), D231–D235. 10.1093/nar/gkab1083.

(2) Slaugenhaupt, S. A.; Blumenfeld, A.; Gill, S. P.; Leyne, M.; Mull, J.; Cuajungco, M. P.; Liebert, C. B.; Chadwick, B.; Idelson, M.; Reznik, L.; Robbins, C.; Makalowska, I.; Brownstein, M.; Krappmann, D.; Scheidereit, C.; Maayan, C.; Axelrod, F. B.; Gusella, J. F. Tissue-Specific Expression of a Splicing Mutation in the IKBKAP Gene Causes Familial Dysautonomia. Am J Hum Genet 2001, 68 (3), 598–605. 10.1086/318810.

(3) Freude, K.; Hoffmann, K.; Jensen, L.-R.; Delatycki, M. B.; des Portes, V.; Moser, B.; Hamel, B.; van Bokhoven, H.; Moraine, C.; Fryns, J.-P.; Chelly, J.; Gécz, J.; Lenzner, S.; Kalscheuer, V. M.; Ropers, H.-H. Mutations in the FTSJ1 Gene Coding for a Novel S-Adenosylmethionine-Binding Protein Cause Nonsyndromic X-Linked Mental Retardation. Am J Hum Genet 2004, 75 (2), 305–309. 10.1086/422507.

(4) Braun, D. A.; Rao, J.; Mollet, G.; Schapiro, D.; Daugeron, M.-C.; Tan, W.; Gribouval, O.; Boyer, O.; Revy, P.; Jobst-Schwan, T.; Schmidt, J. M.; Lawson, J. A.; Schanze, D.; Ashraf, S.; Ullmann, J. F. P.; Hoogstraten, C. A.; Boddaert, N.; Collinet, B.; Martin, G.; Liger, D.; Lovric, S.; Furlano, M.; Guerrera, I. C.; Sanchez-Ferras, O.; Hu, J. F.; Boschat, A.-C.; Sanquer, S.; Menten, B.; Vergult, S.; De Rocker, N.; Airik, M.; Hermle, T.; Shril, S.; Widmeier, E.; Gee, H. Y.; Choi, W.-I.; Sadowski, C. E.; Pabst, W. L.; Warejko, J. K.; Daga, A.; Basta, T.; Matejas, V.; Scharmann, K.; Kienast, S. D.; Behnam, B.; Beeson, B.; Begtrup, A.; Bruce, M.; Ch’ng, G.-S.; Lin, S.-P.; Chang, J.-H.; Chen, C.-H.; Cho, M. T.; Gaffney, P. M.; Gipson, P. E.; Hsu, C.-H.; Kari, J. A.; Ke, Y.-Y.; Kiraly-Borri, C.; Lai, W.-M.; Lemyre, E.; Littlejohn, R. O.; Masri, A.; Moghtaderi, M.; Nakamura, K.; Ozaltin, F.; Praet, M.; Prasad, C.; Prytula, A.; Roeder, E. R.; Rump, P.; Schnur, R. E.; Shiihara, T.; Sinha, M. D.; Soliman, N. A.; Soulami, K.; Sweetser, D. A.; Tsai, W.-H.; Tsai, J.-D.; Topaloglu, R.; Vester, U.; Viskochil, D. H.; Vatanavicharn, N.; Waxler, J. L.; Wierenga, K. J.; Wolf, M. T. F.; Wong, S.-N.; Leidel, S. A.; Truglio, G.; Dedon, P. C.; Poduri, A.; Mane, S.; Lifton, R. P.; Bouchard, M.; Kannu, P.; Chitayat, D.; Magen, D.; Callewaert, B.; van Tilbeurgh, H.; Zenker, M.; Antignac, C.; Hildebrandt, F. Mutations in KEOPS-Complex Genes Cause Nephrotic Syndrome with Primary Microcephaly. Nat Genet 2017, 49 (10), 1529–1538. 10.1038/ng.3933.

(5) Gillis, D.; Krishnamohan, A.; Yaacov, B.; Shaag, A.; Jackman, J. E.; Elpeleg, O. TRMT10A Dysfunction Is Associated with Abnormalities in Glucose Homeostasis, Short Stature and Microcephaly. J Med Genet 2014, 51 (9), 581–586. 10.1136/jmedgenet-2014-102282.

(6) Wei, F.-Y.; Tomizawa, K. Functional Loss of Cdkal1, a Novel tRNA Modification Enzyme, Causes the Development of Type 2 Diabetes. Endocr J 2011, 58 (10), 819–825. 10.1507/endocrj.ej11-0099.

(7) Waszak, S. M.; Robinson, G. W.; Gudenas, B. L.; Smith, K. S.; Forget, A.; Kojic, M.; Garcia-Lopez, J.; Hadley, J.; Hamilton, K. V.; Indersie, E.; Buchhalter, I.; Kerssemakers, J.; Jäger, N.; Sharma, T.; Rausch, T.; Kool, M.; Sturm, D.; Jones, D. T. W.; Vasilyeva, A.; Tatevossian, R. G.; Neale, G.; Lombard, B.; Loew, D.; Nakitandwe, J.; Rusch, M.; Bowers, D. C.; Bendel, A.; Partap, S.; Chintagumpala, M.; Crawford, J.; Gottardo, N. G.; Smith, A.; Dufour, C.; Rutkowski, S.; Eggen, T.; Wesenberg, F.; Kjaerheim, K.; Feychting, M.; Lannering, B.; Schüz, J.; Johansen, C.; Andersen, T. V.; Röösli, M.; Kuehni, C. E.; Grotzer, M.; Remke, M.; Puget, S.; Pajtler, K. W.; Milde, T.; Witt, O.; Ryzhova, M.; Korshunov, A.; Orr, B. A.; Ellison, D. W.; Brugieres, L.; Lichter, P.; Nichols, K. E.; Gajjar, A.; Wainwright, B. J.; Ayrault, O.; Korbel, J. O.; Northcott, P. A.; Pfister, S. M. Germline Elongator Mutations in Sonic Hedgehog Medulloblastoma. Nature 2020, 580 (7803), 396–401. 10.1038/s41586-020-2164-5.

(8) Terao, C.; Suzuki, A.; Momozawa, Y.; Akiyama, M.; Ishigaki, K.; Yamamoto, K.; Matsuda, K.; Murakami, Y.; McCarroll, S. A.; Kubo, M.; Loh, P.-R.; Kamatani, Y. Chromosomal Alterations among Age-Related Haematopoietic Clones in Japan. Nature 2020, 584 (7819), 130–135. 10.1038/s41586-020-2426-2.

(9) Suzuki, T. The Expanding World of tRNA Modifications and Their Disease Relevance. Nat Rev Mol Cell Biol 2021, 22 (6), 375–392. 10.1038/s41580-021-00342-0.

(10) Wang, Y.; Tao, E.-W.; Tan, J.; Gao, Q.-Y.; Chen, Y.-X.; Fang, J.-Y. tRNA Modifications: Insights into Their Role in Human Cancers. Trends Cell Biol 2023, 33 (12), 1035–1048. 10.1016/j.tcb.2023.04.002.

(11) Zhang, W.; Foo, M.; Eren, A. M.; Pan, T. tRNA Modification Dynamics from Individual Organisms to Metaepitranscriptomics of Microbiomes. Mol Cell 2022, 82 (5), 891–906. 10.1016/j.molcel.2021.12.007.

(12) Phizicky, E. M.; Hopper, A. K. The Life and Times of a tRNA. RNA 2023, 29 (7), 898– 957. 10.1261/rna.079620.123.

(13) Torres, A. G.; Piñeyro, D.; Filonava, L.; Stracker, T. H.; Batlle, E.; Ribas de Pouplana, L. A-to-I Editing on tRNAs: Biochemical, Biological and Evolutionary Implications. FEBS Lett 2014, 588 (23), 4279–4286. 10.1016/j.febslet.2014.09.025.

(14) Lorenz, C.; Lünse, C. E.; Mörl, M. tRNA Modifications: Impact on Structure and Thermal Adaptation. Biomolecules 2017, 7 (2). 10.3390/biom7020035.

(15) Vendeix, F. A. P.; Murphy, F. V.; Cantara, W. A.; Leszczyńska, G.; Gustilo, E. M.; Sproat, B.; Malkiewicz, A.; Agris, P. F. Human tRNA(Lys3)(UUU) Is Pre-Structured by Natural Modifications for Cognate and Wobble Codon Binding through Keto-Enol Tautomerism. J Mol Biol 2012, 416 (4), 467–485. 10.1016/j.jmb.2011.12.048.

(16) Grosjean, H.; Westhof, E. An Integrated, Structure- and Energy-Based View of the Genetic Code. Nucleic Acids Res 2016, 44 (17), 8020–8040. 10.1093/nar/gkw608.

(17) Murphy, F. V.; Ramakrishnan, V.; Malkiewicz, A.; Agris, P. F. The Role of Modifications in Codon Discrimination by tRNA(Lys)UUU. Nat Struct Mol Biol 2004, 11 (12), 1186– 1191. 10.1038/nsmb861.

(18) Abbassi, N.-E.-H.; Jaciuk, M.; Scherf, D.; Böhnert, P.; Rau, A.; Hammermeister, A.; Rawski, M.; Indyka, P.; Wazny, G.; Chramiec-Głąbik, A.; Dobosz, D.; Skupien-Rabian, B.; Jankowska, U.; Rappsilber, J.; Schaffrath, R.; Lin, T.-Y.; Glatt, S. Cryo-EM Structures of the Human Elongator Complex at Work. Nat Commun 2024, 15 (1), 4094. 10.1038/s41467-024-48251-y.

(19) Karlsborn, T.; Mahmud, A. K. M. F.; Tükenmez, H.; Byström, A. S. Loss of Ncm5 and Mcm5 Wobble Uridine Side Chains Results in an Altered Metabolic Profile. Metabolomics 2016, 12 (12), 177. 10.1007/s11306-016-1120-8.

(20) Mazauric, M.-H.; Dirick, L.; Purushothaman, S. K.; Björk, G. R.; Lapeyre, B. Trm112p Is a 15-kDa Zinc Finger Protein Essential for the Activity of Two tRNA and One Protein Methyltransferases in Yeast. J Biol Chem 2010, 285 (24), 18505–18515. 10.1074/jbc.M110.113100.

(21) Chen, C.; Huang, B.; Anderson, J. T.; Byström, A. S. Unexpected Accumulation of Ncm(5)U and Ncm(5)S(2) (U) in a Trm9 Mutant Suggests an Additional Step in the Synthesis of Mcm(5)U and Mcm(5)S(2)U. PLoS One 2011, 6 (6), e20783. 10.1371/journal.pone.0020783.

(22) Laxman, S.; Sutter, B. M.; Wu, X.; Kumar, S.; Guo, X.; Trudgian, D. C.; Mirzaei, H.; Tu, B. P. Sulfur Amino Acids Regulate Translational Capacity and Metabolic Homeostasis through Modulation of tRNA Thiolation. Cell 2013, 154 (2), 416–429. 10.1016/j.cell.2013.06.043.

(23) Pabis, M.; Termathe, M.; Ravichandran, K. E.; Kienast, S. D.; Krutyhołowa, R.; Sokołowski, M.; Jankowska, U.; Grudnik, P.; Leidel, S. A.; Glatt, S. Molecular Basis for the Bifunctional Uba4-Urm1 Sulfur-Relay System in tRNA Thiolation and Ubiquitin-like Conjugation. EMBO J 2020, 39 (19), e105087. 10.15252/embj.2020105087.

(24) Rosu, A.; El Hachem, N.; Rapino, F.; Rouault-Pierre, K.; Jorssen, J.; Somja, J.; Ramery, E.; Thiry, M.; Nguyen, L.; Jacquemyn, M.; Daelemans, D.; Adams, C. M.; Bonnet, D.; Chariot, A.; Close, P.; Bureau, F.; Desmet, C. J. Loss of tRNA-Modifying Enzyme Elp3 Activates a P53-Dependent Antitumor Checkpoint in Hematopoiesis. J Exp Med 2021, 218 (3), e20200662. 10.1084/jem.20200662.

(25) Esberg, A.; Huang, B.; Johansson, M. J. O.; Byström, A. S. Elevated Levels of Two tRNA Species Bypass the Requirement for Elongator Complex in Transcription and Exocytosis. Mol Cell 2006, 24 (1), 139–148. 10.1016/j.molcel.2006.07.031.

(26) Nedialkova, D. D.; Leidel, S. A. Optimization of Codon Translation Rates via tRNA Modifications Maintains Proteome Integrity. Cell 2015, 161 (7), 1606–1618. 10.1016/j.cell.2015.05.022.

(27) Dewez, M.; Bauer, F.; Dieu, M.; Raes, M.; Vandenhaute, J.; Hermand, D. The Conserved Wobble Uridine tRNA Thiolase Ctu1-Ctu2 Is Required to Maintain Genome Integrity. Proc Natl Acad Sci U S A 2008, 105 (14), 5459–5464. 10.1073/pnas.0709404105.

(28) Delaunay, S.; Rapino, F.; Tharun, L.; Zhou, Z.; Heukamp, L.; Termathe, M.; Shostak, K.; Klevernic, I.; Florin, A.; Desmecht, H.; Desmet, C. J.; Nguyen, L.; Leidel, S. A.; Willis, E.; Büttner, R.; Chariot, A.; Close, P. Elp3 Links tRNA Modification to IRES-Dependent Translation of LEF1 to Sustain Metastasis in Breast Cancer. J Exp Med 2016, 213 (11), 2503–2523. 10.1084/jem.20160397.

(29) Rapino, F.; Delaunay, S.; Zhou, Z.; Chariot, A.; Close, P. tRNA Modification: Is Cancer Having a Wobble? Trends Cancer 2017, 3 (4), 249–252. 10.1016/j.trecan.2017.02.004.

(30) Pereira, M.; Ribeiro, D. R.; Berg, M.; Tsai, A. P.; Dong, C.; Nho, K.; Kaiser, S.; Moutinho, M.; Soares, A. R. Amyloid Pathology Reduces ELP3 Expression and tRNA Modifications Leading to Impaired Proteostasis. Biochim Biophys Acta Mol Basis Dis 2024, 1870 (1), 166857. 10.1016/j.bbadis.2023.166857.

(31) Suzuki, T.; Suzuki, T. A Complete Landscape of Post-Transcriptional Modifications in Mammalian Mitochondrial tRNAs. Nucleic Acids Res 2014, 42 (11), 7346–7357. 10.1093/nar/gku390.

(32) Chujo, T.; Tomizawa, K. Human Transfer RNA Modopathies: Diseases Caused by Aberrations in Transfer RNA Modifications. FEBS J 2021, 288 (24), 7096–7122. 10.1111/febs.15736.

(33) Boutoual, R.; Jo, H.; Heckenbach, I.; Tiwari, R.; Kasler, H.; Lerner, C. A.; Shah, S.; Schilling, B.; Calvanese, V.; Rardin, M. J.; Scheibye-Knudsen, M.; Verdin, E. A Novel Splice Variant of Elp3/Kat9 Regulates Mitochondrial tRNA Modification and Function. Sci Rep 2022, 12 (1), 14804. 10.1038/s41598-022-18114-x.

(34) Thomas, N. K.; Poodari, V. C.; Jain, M.; Olsen, H. E.; Akeson, M.; Abu-Shumays, R. L. Direct Nanopore Sequencing of Individual Full Length tRNA Strands. ACS Nano 2021, 15 (10), 16642–16653. 10.1021/acsnano.1c06488.

(35) Lucas, M. C.; Pryszcz, L. P.; Medina, R.; Milenkovic, I.; Camacho, N.; Marchand, V.; Motorin, Y.; Ribas de Pouplana, L.; Novoa, E. M. Quantitative Analysis of tRNA Abundance and Modifications by Nanopore RNA Sequencing. Nat Biotechnol 2024, 42 (1), 72–86. 10.1038/s41587-023-01743-6.

(36) Jenjaroenpun, P.; Wongsurawat, T.; Wadley, T. D.; Wassenaar, T. M.; Liu, J.; Dai, Q.; Wanchai, V.; Akel, N. S.; Jamshidi-Parsian, A.; Franco, A. T.; Boysen, G.; Jennings, M. L.; Ussery, D. W.; He, C.; Nookaew, I. Decoding the Epitranscriptional Landscape from Native RNA Sequences. Nucleic Acids Res 2020, 49 (2), e7. 10.1093/nar/gkaa620.

(37) Begik, O.; Lucas, M. C.; Pryszcz, L. P.; Ramirez, J. M.; Medina, R.; Milenkovic, I.; Cruciani, S.; Liu, H.; Vieira, H. G. S.; Sas-Chen, A.; Mattick, J. S.; Schwartz, S.; Novoa, E. M. Quantitative Profiling of Pseudouridylation Dynamics in Native RNAs with Nanopore Sequencing. Nat Biotechnol 2021, 39 (10), 1278–1291. 10.1038/s41587-021-00915-6.

(38) Liu, H.; Begik, O.; Lucas, M. C.; Ramirez, J. M.; Mason, C. E.; Wiener, D.; Schwartz, S.; Mattick, J. S.; Smith, M. A.; Novoa, E. M. Accurate Detection of m6A RNA Modifications in Native RNA Sequences. Nat Commun 2019, 10 (1), 4079. 10.1038/s41467-019-11713-9.

(39) Durant, P. C.; Bajji, A. C.; Sundaram, M.; Kumar, R. K.; Davis, D. R. Structural Effects of Hypermodified Nucleosides in the Escherichia Coli and Human tRNALys Anticodon Loop: The Effect of Nucleosides s2U, mcm5U, mcm5s2U, mnm5s2U, t6A, and ms2t6A. Biochemistry 2005, 44 (22), 8078–8089. 10.1021/bi050343f.

(40) Fu, Y.; Dai, Q.; Zhang, W.; Ren, J.; Pan, T.; He, C. The AlkB Domain of Mammalian ABH8 Catalyzes Hydroxylation of 5-Methoxycarbonylmethyluridine at the Wobble Position of tRNA. Angew Chem Int Ed Engl 2010, 49 (47), 8885–8888. 10.1002/anie.201001242.

(41) Fissekis, J. D.; Sweet, F. Synthesis of 5-Carboxymethyluridine. A Nucleoside from Transfer Ribonucleic Acid. Biochemistry 1970, 9 (16), 3136–3142. 10.1021/bi00818a004.

(42) Bajji, A. C.; Davis, D. R. Synthesis and Biophysical Characterization of tRNA(Lys,3) Anticodon Stem-Loop RNAs Containing the Mcm(5)s(2)U Nucleoside. Org Lett 2000, 2 (24), 3865–3868. 10.1021/ol006605h.

(43) Scaringe, S. A.; Francklyn, C.; Usman, N. Chemical Synthesis of Biologically Active Oligoribonucleotides Using Beta-Cyanoethyl Protected Ribonucleoside Phosphoramidites. Nucleic Acids Res 1990, 18 (18), 5433–5441. 10.1093/nar/18.18.5433.

(44) Wu, T.; Ogilvie, K. K.; Pon, R. T. Prevention of Chain Cleavage in the Chemical Synthesis of 2’-Silylated Oligoribonucleotides. Nucleic Acids Res 1989, 17 (9), 3501–3517. 10.1093/nar/17.9.3501.

(45) Stawinski, J.; Strömberg, R.; Thelin, M.; Westman, E. Studies on the T-Butyldimethylsilyl Group as 2’-O-Protection in Oligoribonucleotide Synthesis via the H-Phosphonate Approach. Nucleic Acids Res 1988, 16 (19), 9285–9298. 10.1093/nar/16.19.9285.

(46) Eshete, M.; Marchbank, M. T.; Deutscher, S. L.; Sproat, B.; Leszczynska, G.; Malkiewicz, A.; Agris, P. F. Specificity of Phage Display Selected Peptides for Modified Anticodon Stem and Loop Domains of tRNA. Protein J 2007, 26 (1), 61–73. 10.1007/s10930-006-9046-z.

(47) Saville, L.; Wu, L.; Habtewold, J.; Cheng, Y.; Gollen, B.; Mitchell, L.; Stuart-Edwards, M.; Haight, T.; Mohajerani, M.; Zovoilis, A. NERD-Seq: A Novel Approach of Nanopore Direct RNA Sequencing That Expands Representation of Non-Coding RNAs. Genome Biol 2024, 25 (1), 233. 10.1186/s13059-024-03375-8.

(48) Kholod, N.; Vassilenko, K.; Shlyapnikov, M.; Ksenzenko, V.; Kisselev, L. Preparation of Active tRNA Gene Transcripts Devoid of 3’-Extended Products and Dimers. Nucleic Acids Res 1998, 26 (10), 2500–2501. 10.1093/nar/26.10.2500.

(49) Stephenson, W.; Razaghi, R.; Busan, S.; Weeks, K. M.; Timp, W.; Smibert, P. Direct Detection of RNA Modifications and Structure Using Single-Molecule Nanopore Sequencing. Cell Genom 2022, 2 (2), 100097. 10.1016/j.xgen.2022.100097.

(50) Leger, A.; Amaral, P. P.; Pandolfini, L.; Capitanchik, C.; Capraro, F.; Miano, V.; Migliori, V.; Toolan-Kerr, P.; Sideri, T.; Enright, A. J.; Tzelepis, K.; van Werven, F. J.; Luscombe, N. M.; Barbieri, I.; Ule, J.; Fitzgerald, T.; Birney, E.; Leonardi, T.; Kouzarides, T. RNA Modifications Detection by Comparative Nanopore Direct RNA Sequencing. Nat Commun 2021, 12 (1), 7198. 10.1038/s41467-021-27393-3.

(51) Parker, M. T.; Knop, K.; Sherwood, A. V.; Schurch, N. J.; Mackinnon, K.; Gould, P. D.; Hall, A. J.; Barton, G. J.; Simpson, G. G. Nanopore Direct RNA Sequencing Maps the Complexity of Arabidopsis mRNA Processing and m6A Modification. Elife 2020, 9, e49658. 10.7554/eLife.49658.

(52) Pratanwanich, P. N.; Yao, F.; Chen, Y.; Koh, C. W. Q.; Wan, Y. K.; Hendra, C.; Poon, P.; Goh, Y. T.; Yap, P. M. L.; Chooi, J. Y.; Chng, W. J.; Ng, S. B.; Thiery, A.; Goh, W. S. S.; Göke, J. Identification of Differential RNA Modifications from Nanopore Direct RNA Sequencing with xPore. Nat Biotechnol 2021, 39 (11), 1394–1402. 10.1038/s41587-021-00949-w.

(53) Price, A. M.; Hayer, K. E.; McIntyre, A. B. R.; Gokhale, N. S.; Abebe, J. S.; Della Fera, A. N.; Mason, C. E.; Horner, S. M.; Wilson, A. C.; Depledge, D. P.; Weitzman, M. D. Direct RNA Sequencing Reveals m6A Modifications on Adenovirus RNA Are Necessary for Efficient Splicing. Nat Commun 2020, 11 (1), 6016. 10.1038/s41467-020-19787-6.

(54) White, L. K.; Dobson, K.; Del Pozo, S.; Bilodeaux, J. M.; Andersen, S. E.; Baldwin, A.; Barrington, C.; Körtel, N.; Martinez-Seidel, F.; Strugar, S. M.; Watt, K. E. N.; Mukherjee, N.; Hesselberth, J. R. Comparative Analysis of 43 Distinct RNA Modifications by Nanopore tRNA Sequencing. bioRxiv 2024, 2024.07.23.604651. 10.1101/2024.07.23.604651.

(55) Guo, Z.; Ni, Y.; Tan, L.; Shao, Y.; Ye, L.; Chen, S.; Li, R. Nanopore Current Events Magnifier (nanoCEM): A Novel Tool for Visualizing Current Events at Modification Sites of Nanopore Sequencing. NAR Genom Bioinform 2024, 6 (2), lqae052. 10.1093/nargab/lqae052.

(56) Lang, K.; Micura, R. The Preparation of Site-Specifically Modified Riboswitch Domains as an Example for Enzymatic Ligation of Chemically Synthesized RNA Fragments. Nat Protoc 2008, 3 (9), 1457–1466. 10.1038/nprot.2008.135.

(57) Su, T.; Hollas, M. A. R.; Fellers, R. T.; Kelleher, N. L. Identification of Splice Variants and Isoforms in Transcriptomics and Proteomics. Annu Rev Biomed Data Sci 2023, 6, 357–376. 10.1146/annurev-biodatasci-020722-044021.

(58) Boccaletto, P.; Bagiński, B. MODOMICS: An Operational Guide to the Use of the RNA Modification Pathways Database. Methods Mol Biol 2021, 2284, 481–505. 10.1007/978-1-0716-1307-8_26.

(59) Li, H. Minimap2: Pairwise Alignment for Nucleotide Sequences. Bioinformatics 2018, 34 (18), 3094–3100. 10.1093/bioinformatics/bty191.

(60) Danecek, P.; Bonfield, J. K.; Liddle, J.; Marshall, J.; Ohan, V.; Pollard, M. O.; Whitwham, A.; Keane, T.; McCarthy, S. A.; Davies, R. M.; Li, H. Twelve Years of SAMtools and BCFtools. Gigascience 2021, 10 (2), giab008. 10.1093/gigascience/giab008.

(61) Robinson, J. T.; Thorvaldsdóttir, H.; Winckler, W.; Guttman, M.; Lander, E. S.; Getz, G.; Mesirov, J. P. Integrative Genomics Viewer. Nat Biotechnol 2011, 29 (1), 24–26. 10.1038/nbt.1754.

(62) Thorvaldsdóttir, H.; Robinson, J. T.; Mesirov, J. P. Integrative Genomics Viewer (IGV): High-Performance Genomics Data Visualization and Exploration. Brief Bioinform 2013, 14 (2), 178–192. 10.1093/bib/bbs017.

(63) Robinson, J. T.; Thorvaldsdóttir, H.; Wenger, A. M.; Zehir, A.; Mesirov, J. P. Variant Review with the Integrative Genomics Viewer. Cancer Res 2017, 77 (21), e31–e34. 10.1158/0008-5472.CAN-17-0337.

(64) Robinson, J. T.; Thorvaldsdottir, H.; Turner, D.; Mesirov, J. P. Igv.Js: An Embeddable JavaScript Implementation of the Integrative Genomics Viewer (IGV). Bioinformatics 2023, 39 (1), btac830. 10.1093/bioinformatics/btac830.

