## Supporting information for "Direct RNA Oxford Nanopore sequencing distinguishes between modifications in tRNA at the U_34_ position"

‡ first co-authors

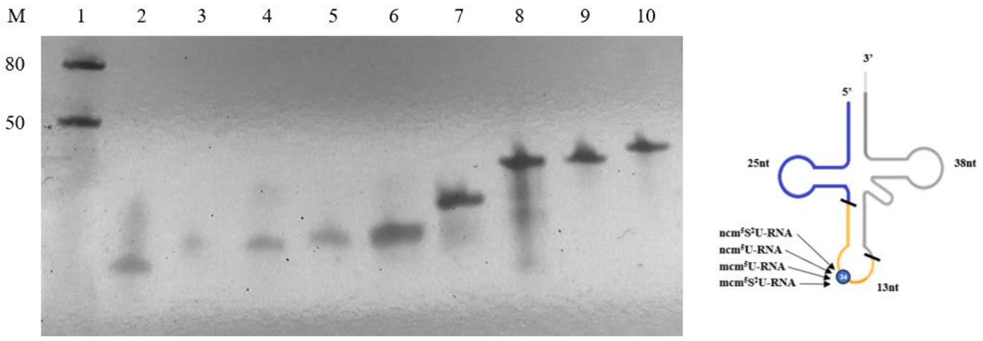


**Figure S1 RNA and DNA oligos used in splint ligation reactions.** line 1: Low range ssRNA ladder. List of the oligonucleotides: see below.

| Line No. | Oligo name | Sequence 5'-3' |
|---|---|---|
| 2 | 13nt-RNA | [Phos]GUUCGGCUUUUAA |
| 3 | 13nt- RNA $ncm^5U$ | GUUCGGCUncm$^5$UUUAA |
| 4 | 13nt- RNA $ncm^5S^2U$ | GUUCGGCUncm$^5$S$^2$UUUAA |
| 5 | 13nt-RNA $mcm^5U$ | GUUCGGCUmcm$^5$S$^2$UUUAA |
| 6 | 13nt-RNA $mcm^5S^2U$ | GUUCGGCUmcm$^5$S$^2$UUUAA |
| 7 | 25nt-RNA | [Phos]UCCUUGUUAGCUCAGUUGGUAGAGC |
| 8 | 38nt-RNA | [Phos]CCGAAAUGUCAGGGGUUCGAGCCCCCUAUGAGGAGCCA |

| | | |
|---|---|---|
| **9** | **Splint DNA no.2a** | TAGGGGGCTCGAACCCCTGACATTTCGGTTAAAAGCCGAACGCTCT |
| **10** | **Splint DNA no. 1** | GGTTAAAAGCCGAACGCTCTACCAACTGAGCTAACAAGGA |

**Table S1 Sequences of RNA and DNA oligos displayed on Fig. S1.**

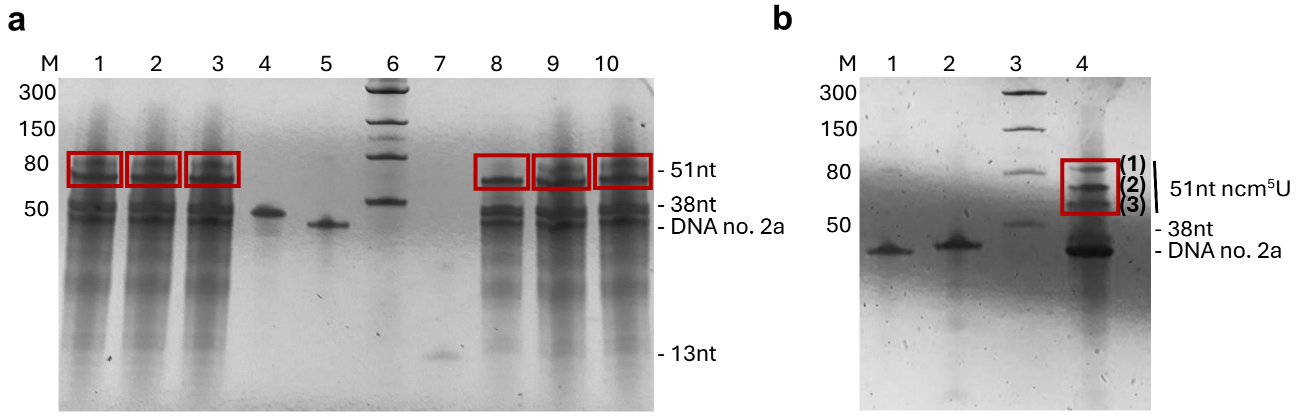


**Figure S2 The ligation products migration pattern on UREA-PAGE before reaction optimisation**. (a) lines 1-3 and 8-10: ligation mixtures containing 13nt-RNA unmodified with 38nt-RNA.; line 4: 38nt-RNA; line 5: DNA no.2a, line 6: ssRNA MW leader, line 7: 13nt-RNA (b) line 1: 38nt-RNA; line 2: DNA no.2a; line 3: Low Range ssRNA ladder (NEB); line 4: ligation mixtures of13nt-RNA ncm$^5$U with 38nt-RNA. The unmodified and modified tRNA$^{Lys}$ ligation product bands are highlighted in red and identified by LC-MS (Fig 3b, Fig3c).

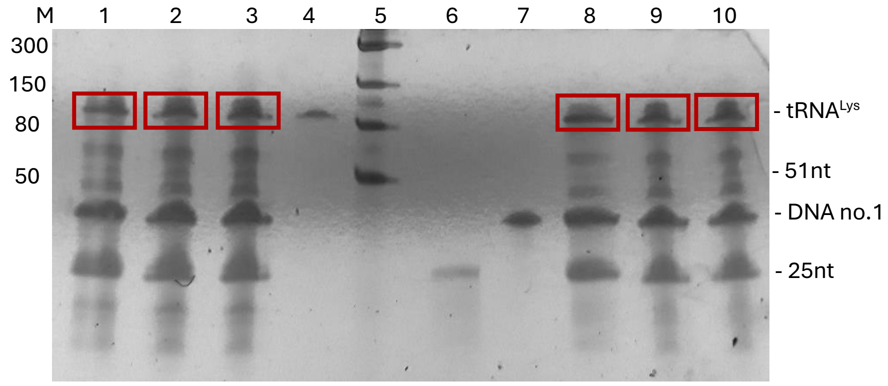


**Figure S3 UREA-PAGE of the splint ligation reaction mixtures without DNase I treatment.** ligation II: lines 1-3: unmodified tRNA$^{Lys}$ ligation product; line 4: control tRNA$^{Lys}$ ncm$^5$U; line 5: Low Range ssRNA ladder; line 6: 25nt-RNA; line 6: DNA no.1; lines 8-10: unmodified tRNA$^{Lys}$. The unmodified full tRNA$^{Lys}$ ligation product bands are highlighted in red.

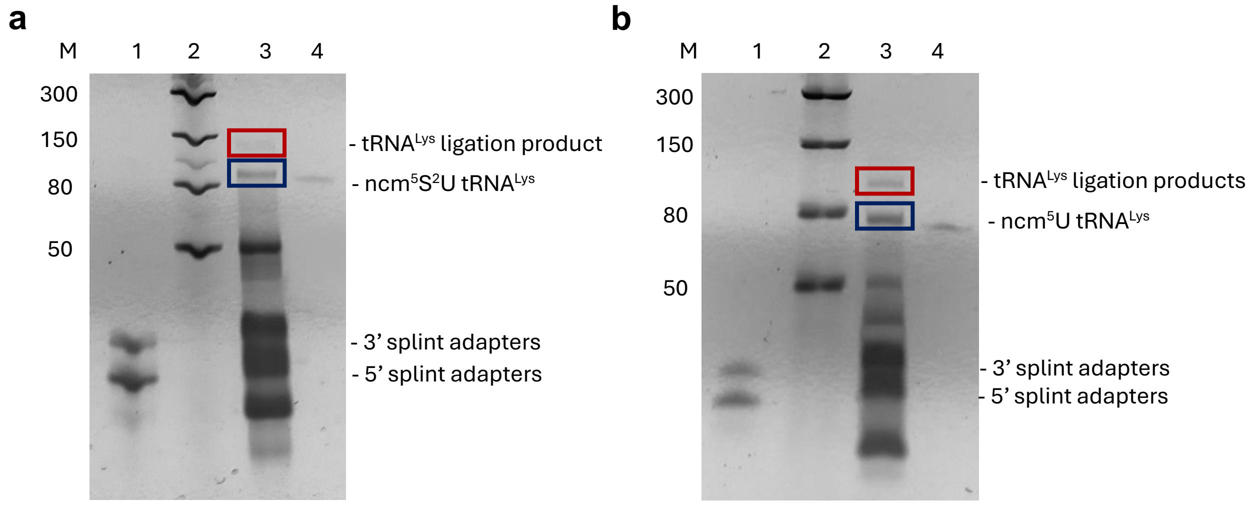


**Figure S4 UREA-PAGE of ligation reaction mixtures of tRNA$^{Lys}$ with adapters.** (a) line 1: 3’ splint oligo, 5’ splint oligo; line 2: Low Range ssRNA ladder; line 3: tRNA$^{Lys}$ ncm$^5$S$^2$U; line 2: ligation products of tRNALys ncm$^5$S$^2$U with adapters; line 3:; line 4: 5’ splint and 3’ splint adaptors; line 5: ligation products of unmodified tRNA$^{Lys}$ with adapters; line6: unmodified tRNA$^{Lys}$ (b) line 1: tRNA$^{Lys}$ ncm$^5$U; line 2: ligation products of tRNA$^{Lys}$ ncm$^5$U with adapters; line 3: Low Range ssRNA ladder; line 4: 5’ splint and 3’ splint adapters; line 5: ligation products of tRNALys mcm$^5$S$^2$U; line 6: tRNA$^{Lys}$ mcm$^5$S$^2$U; line 7: ligation products of tRNA$^{Lys}$ mcm$^5$U; line 8: tRNA$^{Lys}$ mcm$^5$U. The ligation products are highlighted in red.

| | Estimated bases [Mb] | Reads generated [k] | Estimated N50 [kb] | Total data produced (pass / fail) | Time [h] |
|---|---|---|---|---|---|
| control | 146.54 | 200.6 | 3.99 | 7.83 GB | 60 |
| ncm$^5$U$_{34}$ | 17.57 | 8.43 | 5.12 | 856.6 MB | 42 |
| ncm$^5$S$^2$U$_{34}$ | 13.2 | 4.98 | 9.37 | 612.9 MB | 72 |

**Table S2**. **Basic sequencing process statistics from a run report for a sequencing run.**

| | Number of RNA reads | | | | Sum of bases [MB] | | | |
|---|---|---|---|---|---|---|---|---|
| | dorado hac | dorado fast | guppy pass | guppy fail | dorado hac | dorado fast | guppy pass | guppy fail |
| control | 200466 | 200529 | 88868 | 111729 | 58.638 | 55.032 | 8.753 | 35.797 |
| $ncm^5U_{34}$ | 8384 | 8408 | 591 | 7834 | 5.354 | 4.375 | 0.121 | 3.811 |
| $ncm^5S^2U_{34}$ | 9906 | 9946 | 374 | 4605 | 8.673 | 6.870 | 0.297 | 2.884 |

**Table S3**. **Number of RNA reads, and sum of bases included in all RNA reads obtained from different basecallers.**

| | Number of RNA reads | | | | Sum of bases [MB] | | | |
|---|---|---|---|---|---|---|---|---|
| | dorado hac | dorado fast | guppy pass | guppy fail | dorado hac | dorado fast | guppy pass | guppy fail |
| control | 92810 | 107922 | 88393 | 9186 | 13.340 | 16.022 | 7.667 | 1.025 |
| $ncm^5U_{34}$ | 433 | 557 | 528 | 53 | 0.053 | 0.073 | 0.038 | 0.004 |
| $ncm^5S^2U_{34}$ | 360 | 726 | 312 | 48 | 0.052 | 0.108 | 0.028 | 0.005 |

**Table S4**. **Number of RNA reads and their sum of bases after discarding reads longer than 200 bp and with a mean base call accuracy lower than 80%.**

| | Mapping parameters | Mapped reads (#) | Mapped reads (%) | Uniquely mapped reads (#) | Uniquely mapped reads (%) | Antisense (#) | Antisense (%) |
|---|---|---|---|---|---|---|---|
| control | minimap2 -ax map-ont -k15 | 9286 | 10.01 | 9286 | 100.00 | 0 | 0.00 |
| | bwa mem -W13 -k6 - | 65548 | 70.63 | 65546 | 100.00 | 0 | 0.00 |

| | | | | | | | |
|---|---|---|---|---|---|---|---|
| | xont2d -T30 | | | | | | |
| | bwa mem -W13 -k6 -xont2d -T20 | 78739 | 84.84 | 78545 | 99.75 | 85 | 0.11 |
| | bwa mem -W13 -k6 -xont2d -T10 | 88246 | 95.08 | 86010 | 97.47 | 1065 | 1.24 |
| | bwa mem -W9 -k5 -xont2d -T10 | 90320 | 97.72 | 87020 | 96.35 | 1030 | 1.18 |
| | bwa sw -z10 -a2 -b1 -q2 -r1 | 55706 | 51.09 | 28147 | 50.53 | 5830 | 20.71 |
| $ncm^5U_{34}$ | minimap2 -ax map-ont -k15 | 6 | 1.39 | 6 | 100.00 | 0 | 0.00 |
| | bwa mem -W13 -k6 -xont2d -T30 | 169 | 39.03 | 169 | 100.00 | 0 | 0.00 |
| | bwa mem -W13 -k6 -xont2d -T20 | 271 | 62.59 | 263 | 97.05 | 8 | 3.04 |
| | bwa mem -W13 -k6 -xont2d -T10 | 365 | 84.30 | 317 | 86.85 | 35 | 11.04 |
| | bwa mem -W9 -k5 -xont2d -T10 | 388 | 89.61 | 319 | 82.22 | 41 | 12.85 |

| | | | | | | | |
|---|---|---|---|---|---|---|---|
| | bwa sw -z10 -a2 -b1 -q2 -r1 | 92 | 20.40 | 45 | 48.91 | 11 | 24.44 |
| ncm$^5$S$^2$U$_{34}$ | minimap2 -ax map-ont -k15 | 24 | 6.67 | 24 | 100.00 | 0 | 0.00 |
| | bwa mem -W13 -k6 -xont2d -T30 | 220 | 61.11 | 220 | 100.00 | 0 | 0.00 |
| | bwa mem -W13 -k6 -xont2d -T20 | 276 | 76.67 | 276 | 100.00 | 0 | 0.00 |
| | bwa mem -W13 -k6 -xont2d -T10 | 323 | 89.72 | 310 | 95.98 | 9 | 2.90 |
| | bwa mem -W9 -k5 -xont2d -T10 | 337 | 93.61 | 311 | 92.28 | 10 | 3.22 |
| | bwa sw -z10 -a2 -b1 -q2 -r1 | 225 | 53.32 | 104 | 46.22 | 22 | 21.15 |

**Table S5. Mapping statistics of reads from control, ncm$^5$U$_{34}$ and ncm$^5$S$^2$U$_{34}$ samples from dorado basecaller with rna002_70bps_hac@v3 model using different applications and parameters.**

| | Mapping parameters | Mapped reads (#) | Mapped reads (%) | Uniquely mapped reads (#) | Uniquely mapped reads (%) | Antisense (#) | Antisense (%) |
|---|---|---|---|---|---|---|---|
| control | minimap2 -ax map-ont -k15 | 8009 | 7.42 | 8009 | 100.00 | 0 | 0.00 |
| | bwa mem -W13 -k6 -xont2d -T30 | 84269 | 78.08 | 84264 | 99.99 | 0 | 0.00 |
| | bwa mem -W13 -k6 -xont2d -T20 | 99245 | 91.96 | 99122 | 99.88 | 59 | 0.06 |
| | bwa mem -W13 -k6 -xont2d -T10 | 106230 | 98.43 | 104728 | 98.59 | 739 | 0.71 |
| | bwa mem -W9 -k5 -xont2d -T10 | 107181 | 99.31 | 105208 | 98.16 | 562 | 0.53 |
| | bwa sw -z10 -a2 -b1 -q2 -r1 | 70931 | 54.59 | 35350 | 49.84 | 6111 | 17.29 |
| $ncm^5U_{34}$ | minimap2 -ax map-ont -k15 | 16 | 2.87 | 16 | 100.00 | 0 | 0.00 |
| | bwa mem -W13 -k6 -xont2d -T30 | 314 | 56.37 | 314 | 100.00 | 0 | 0.00 |
| | bwa mem -W13 -k6 - | 412 | 73.97 | 407 | 98.79 | 5 | 1.23 |

| | | | | | | | |
|---|---|---|---|---|---|---|---|
| | xont2d - T20 | | | | | | |
| | bwa mem - W13 -k6 - xont2d - T10 | 527 | 94.61 | 453 | 85.96 | 46 | 10.15 |
| | bwa mem - W9 -k5 - xont2d - T10 | 537 | 96.41 | 456 | 84.92 | 49 | 10.75 |
| | bwa sw - z10 -a2 -b1 -q2 -r1 | 172 | 28.52 | 88 | 51.16 | 15 | 17.05 |
| $ncm^5S^2U_{34}$ | minimap2 - ax map-ont -k15 | 30 | 4.13 | 30 | 100.00 | 0 | 0.00 |
| | bwa mem - W13 -k6 - xont2d - T30 | 558 | 76.86 | 558 | 100.00 | 0 | 0.00 |
| | bwa mem - W13 -k6 - xont2d - T20 | 652 | 89.81 | 648 | 99.39 | 0 | 0.00 |
| | bwa mem - W13 -k6 - xont2d - T10 | 696 | 95.87 | 684 | 98.28 | 4 | 0.58 |
| | bwa mem - W9 -k5 - xont2d - T10 | 702 | 96.69 | 684 | 97.44 | 4 | 0.58 |
| | bwa sw - z10 -a2 -b1 -q2 -r1 | 472 | 54.50 | 250 | 52.97 | 32 | 12,8 |

**Table S6. Mapping statistics of reads from control, ncm$^5$U$_{34}$ and ncm$^5$S$^2$U$_{34}$ samples from dorado basecaller with rna002_70bps_fast@v3 model using different applications and parameters.**

| | Mapping parameters | Mapped reads (#) | Mapped reads (%) | Uniquely mapped reads (#) | Uniquely mapped reads (%) | Antisense (#) | Antisense (%) |
|---|---|---|---|---|---|---|---|
| control | minimap2 -ax map-ont -k15 | 5438 | 6.15 | 5438 | 100.00 | 0 | 0.00 |
| | bwa mem -W13 -k6 -xont2d -T30 | 75042 | 84.90 | 75039 | 100.00 | 0 | 0.00 |
| | bwa mem -W13 -k6 -xont2d -T20 | 85311 | 96.51 | 85262 | 99.94 | 27 | 0.03 |
| | bwa mem -W13 -k6 -xont2d -T10 | 87870 | 99.41 | 87529 | 99.61 | 194 | 0.22 |
| | bwa mem -W9 -k5 -xont2d -T10 | 88135 | 99.71 | 87729 | 99.54 | 157 | 0.18 |
| | bwa sw -z10 -a2 -b1 -q2 -r1 | 65000 | 70.76 | 60005 | 92.32 | 1563 | 2.60 |
| ncm$^5$U$_{34}$ | minimap2 -ax map-ont -k15 | 13 | 2.46 | 13 | 100.00 | 0 | 0.00 |
| | bwa mem -W13 -k6 - | 386 | 73.11 | 386 | 100.00 | 0 | 0.00 |

| | | | | | | | |
|---|---|---|---|---|---|---|---|
| | xont2d -T30 | | | | | | |
| | bwa mem -W13 -k6 -xont2d -T20 | 459 | 86.93 | 454 | 98.91 | 5 | 1.10 |
| | bwa mem -W13 -k6 -xont2d -T10 | 501 | 94.89 | 473 | 94.41 | 23 | 4.86 |
| | bwa mem -W9 -k5 -xont2d -T10 | 504 | 95.45 | 471 | 93.45 | 22 | 4.67 |
| | bwa sw -z10 -a2 -b1 -q2 -r1 | 357 | 67.11 | 349 | 97.76 | 0 | 0.00 |
| $ncm^5S^2U_{34}$ | minimap2 -ax map-ont -k15 | 20 | 6.41 | 20 | 100.00 | 0 | 0.00 |
| | bwa mem -W13 -k6 -xont2d -T30 | 256 | 82.05 | 256 | 100.00 | 0 | 0.00 |
| | bwa mem -W13 -k6 -xont2d -T20 | 289 | 92.63 | 288 | 99.65 | 1 | 0.35 |
| | bwa mem -W13 -k6 -xont2d -T10 | 304 | 97.44 | 299 | 98.36 | 4 | 1.34 |
| | bwa mem -W9 -k5 -xont2d -T10 | 305 | 97.76 | 298 | 97.70 | 3 | 1.01 |

| | | | | | | | |
|---|---|---|---|---|---|---|---|
| | bwa sw -z10 -a2 -b1 -q2 -r1 | 244 | 73.72 | 217 | 88.93 | 10 | 4.61 |

**Table S7**. **Mapping statistics of reads from control, ncm$^5$U$_{34}$ and ncm$^5$S$^2$U$_{34}$ samples from guppy basecaller with guppy_rna_r9.4.1_70bps_hac model (pass subset) using different applications and parameters.**

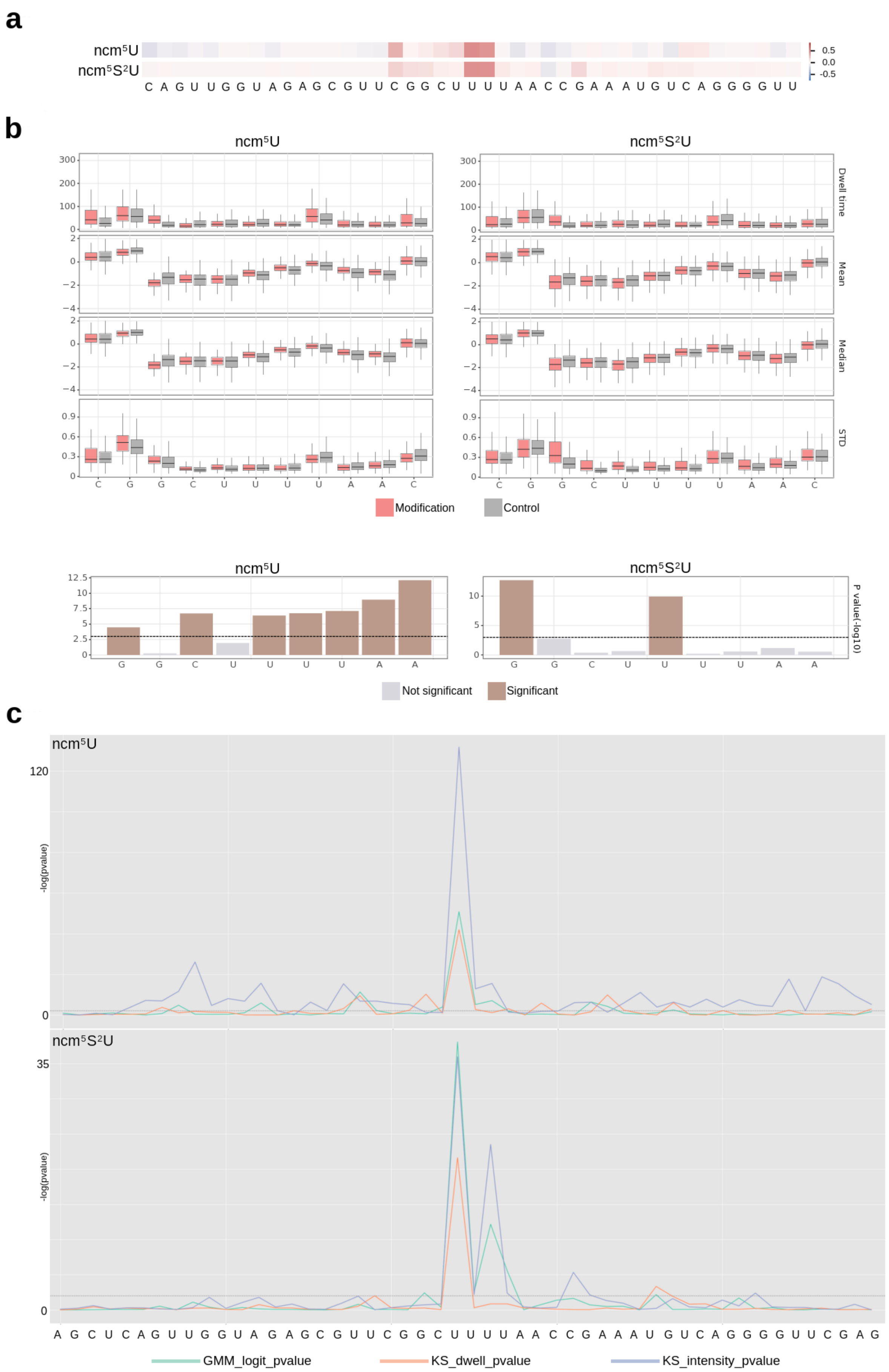
a
ncm⁵U
ncm⁵S²U
0.5
0.0
-0.5
C A G U U G G U A G A G C G U U C G G C U U U U A A C C G A A A U G U C A G G G G U U
b
ncm⁵U
ncm⁵S²U
Dwell time
Mean
Median
STD
C G G C U U U U A A C
Modification
Control
ncm⁵U
ncm⁵S²U
P value(-log10)
G G C U U U U A A
Not significant
Significant
c
ncm⁵U
-log(pvalue)
120
0
ncm⁵S²U
35
0
A G C U C A G U U G G U A G A G C G U U C G G C U U U U A A C C G A A A U G U C A G G G G U U C G A G
GMM_logit_pvalue
KS_dwell_pvalue
KS_intensity_pvalue

**Figure S5. Detection of $ncm^5U_{34}$ and $ncm^5S^2U_{34}$ $tRNA^{Lys}$ synthetic modifications using Nano-tRNAseq (a), nanoCEM (b) and Nanocompore (c) tools - dorado basecaller with rna002_70bps_fast@v3 model.**

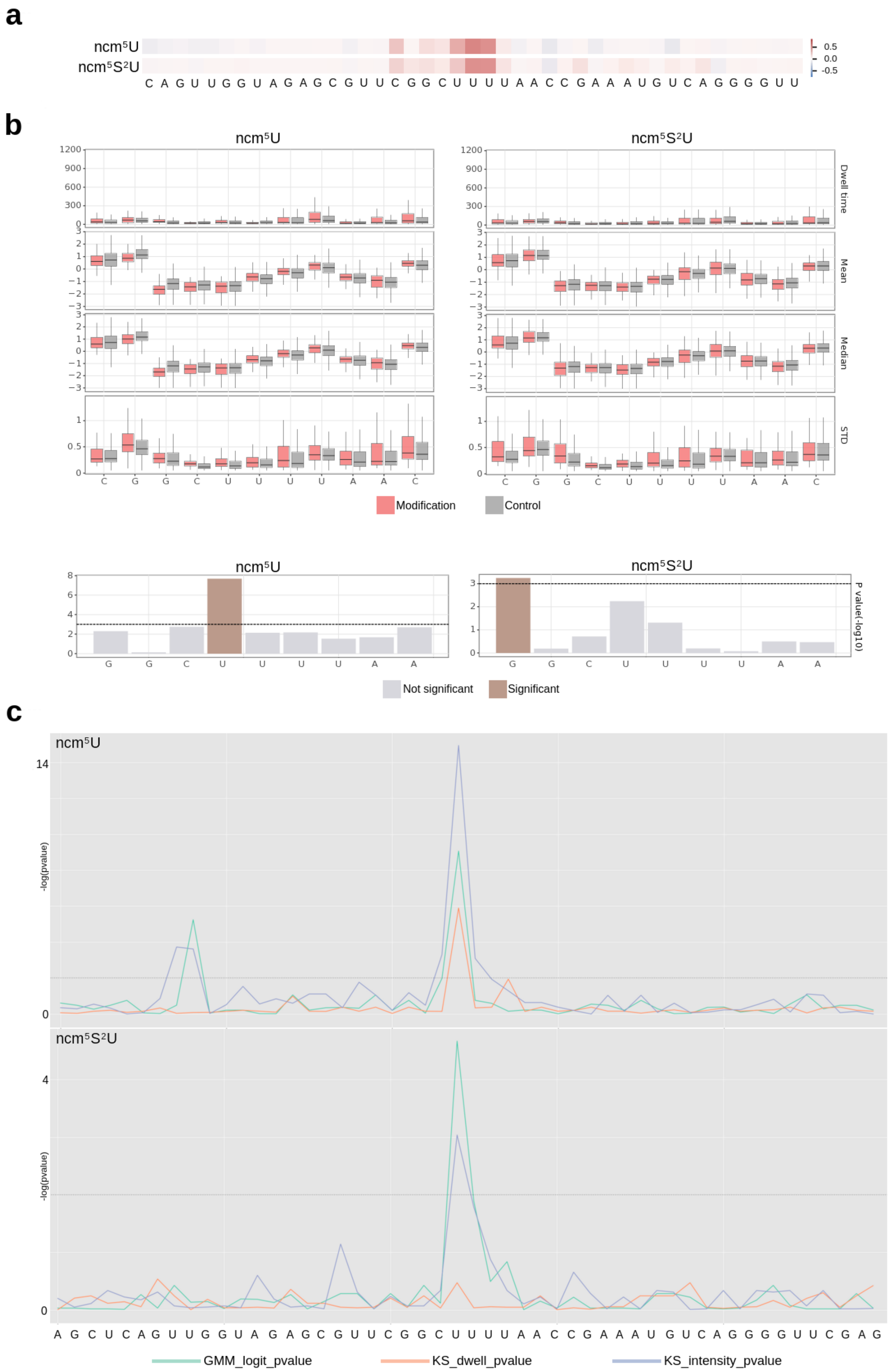

a
ncm5U
ncm5S2U
0.5
0.0
-0.5
C A G U U G G U A G A G C G U U C G G C U U U U A A C C G A A A U G U C A G G G G U U
b
ncm5U
ncm5S2U
Dwell time
Mean
Median
STD
Modification
Control
P value(-log10)
Not significant
Significant
c
ncm5U
ncm5S2U
-log(pvalue)
A G C U C A G U U G G U A G A G C G U U C G G C U U U U A A C C G A A A U G U C A G G G G U U C G A G
GMM_logit_pvalue
KS_dwell_pvalue
KS_intensity_pvalue

**Figure S6. Detection of $ncm^5U_{34}$ and $ncm^5S^2U_{34}$ in $tRNA^{Lys}$ synthetic modifications using Nano-tRNAseq (a), nanoCEM (b) and Nanocompore (c) tools - guppy basecaller with guppy_rna_r9.4.1_70bps_hac model (pass subset).**

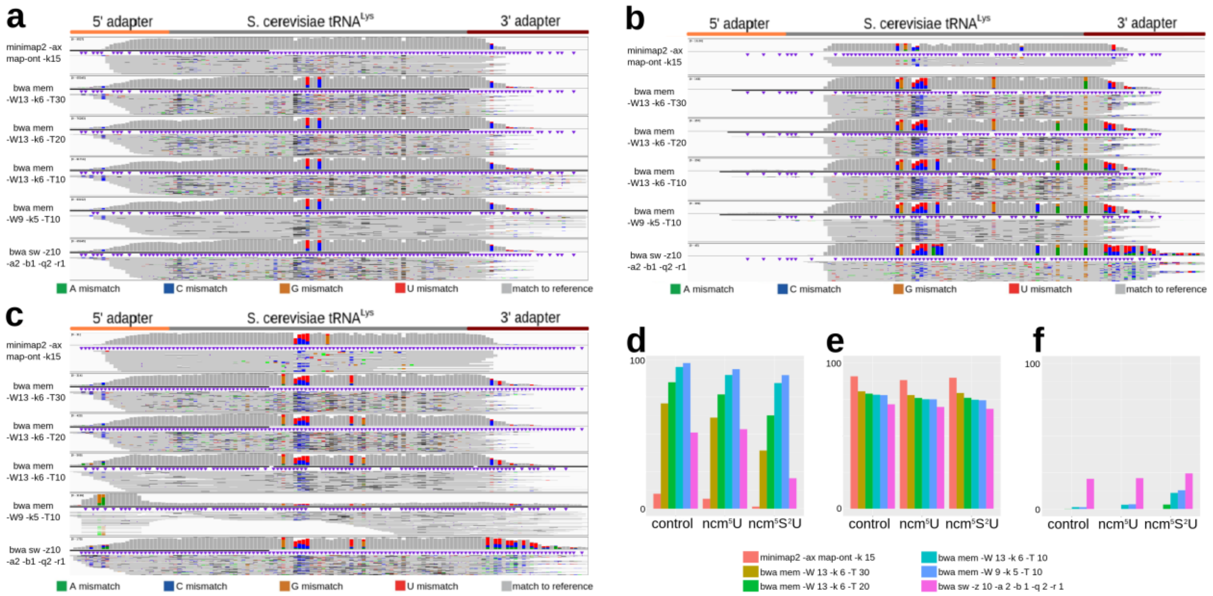


**Figure. S7. Influence of the mapping algorithm and its parameters on the mapping results - dorado basecaller with rna002_70bps_hac@v3 model.** (a), (b) and (c) present the effect of the mapping algorithm and its parameters on the base pair error distribution for the control $tRNA^{Lys}$, $ncm^5U_{34}$-$tRNA^{Lys}$ and $ncm^5S^2U_{34}$-$tRNA^{Lys}$, respectively. (d) Percentage of reads mapped to the reference sequence. (e) Average mapping alignment identity. (f) Percentage of mismapped reads - reads mapped to antisense strand.

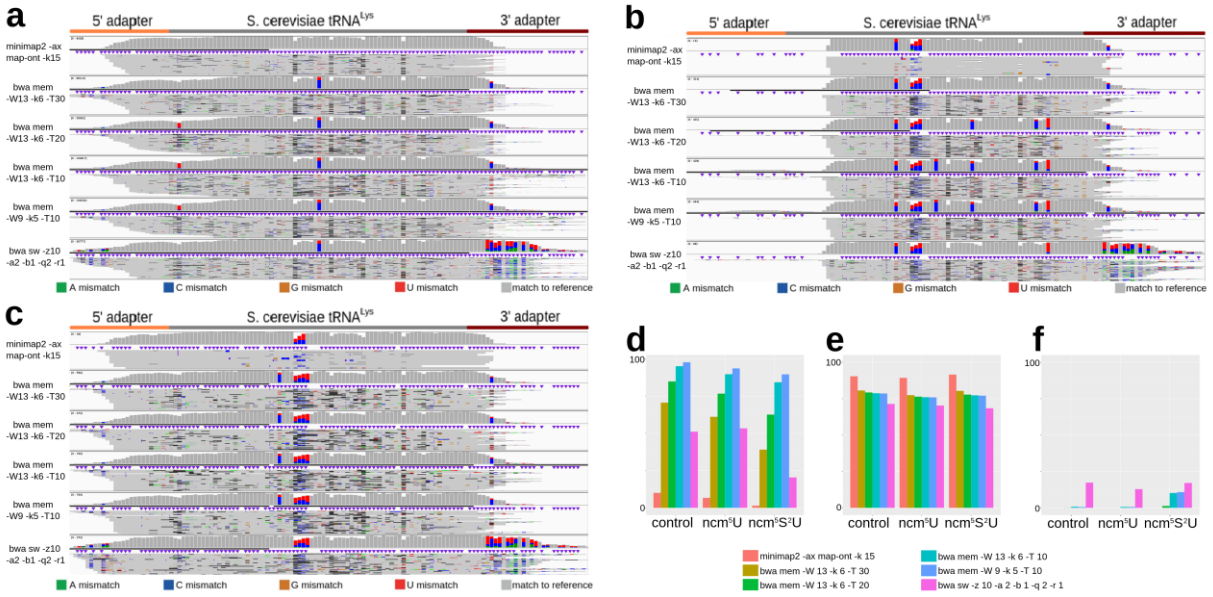


**Fig. S8. Influence of the mapping algorithm and its parameters on the mapping results - dorado basecaller with rna002_70bps_fast@v3 model.** (a), (b) and (c) present the effect of the mapping algorithm and its parameters on the base pair error distribution for the control tRNA$^{Lys}$, ncm$^5$U$_{34}$-tRNA$^{Lys}$ and ncm$^5$S$^2$U$_{34}$-tRNA$^{Lys}$, respectively. (d) Percentage of reads mapped to the reference sequence. (e) Average mapping alignment identity. (f) Percentage of mismapped reads - reads mapped to antisense strand.

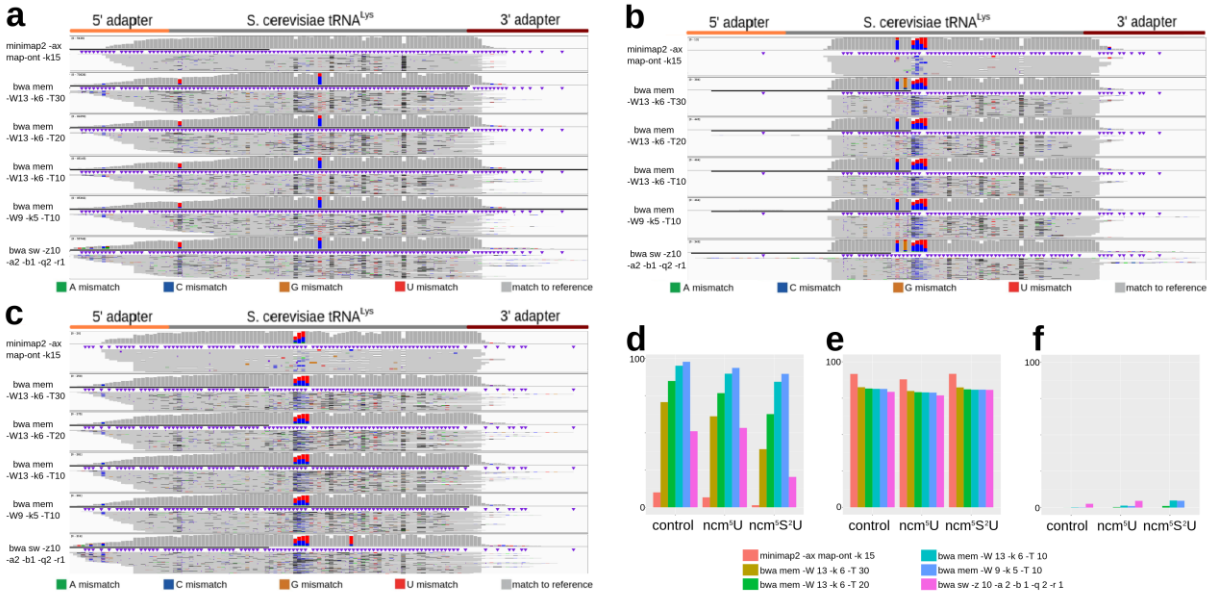


**Figure S9. Influence of the mapping algorithm and its parameters on the mapping results - guppy basecaller with guppy_rna_r9.4.1_70bps_hac model (pass subset).** (a), (b) and (c)

present the effect of the mapping algorithm and its parameters on the base pair error distribution for the control $tRNA^{Lys}$, $ncm^5U_{34}$-$tRNA^{Lys}$ and $ncm^5S^2U_{34}$-$tRNA^{Lys}$, respectively. (d) Percentage of reads mapped to the reference sequence. (e) Average mapping alignment identity. (f) Percentage of mismapped reads - reads mapped to antisense strand.
